# Benchmarking the Performance of a Commercial Electrostatic Linear Ion Trap (ELIT) Charge Detection Mass Spectrometer For Higher-Order Antibody Aggregate Analysis

**DOI:** 10.64898/2026.10.08.757778

**Authors:** Shang-Yin Wu, Xing-Miao Guo, Jia-Yi Lv, Ying-Zi Li, Lei Gu, Zachary Tian, Yong-Sheng Xiao, Emily Christofi, Anisha Haris, Haoyun Lu, Rui Wang, Huan-Fang Xie, Hongwen Li, Yuanjiang Pan, Mowei Zhou, Wei-Jie Fang

**Affiliations:** Institute of Drug Metabolism and Pharmaceutical Analysis, School of Pharmacy, Zhejiang University, Hangzhou, 310058, China; Taizhou Institute of Zhejiang University, Taizhou, 318000, China; Department of Chemistry, Zhejiang University, Hangzhou, 310058, China; WuXi Biologies Co., Ltd, Jiangsu, 214092, China; Waters Corporation, Wilmslow, SK9 4AX, UK; Institute of Fundamental and Transdisciplinary Research, Zhejiang University, Hangzhou, 310058, China; Hangzhou Zhongmei Huadong pharmaceutical Co., Ltd, Hangzhou, 310058, China

## Abstract

Higher-order antibody aggregates remain challenging to characterize using conventional mass spectrometry because of their low abundance, broad mass distributions, and extensive heterogeneity. Here, the practical performance of a commercial electrostatic linear ion trap charge detection mass spectrometer (Xevo^TM^ CDMS) was evaluated for therapeutic antibody analysis. The platform provided reproducible mass and charge measurements, with repeatability and intermediate-precision RSDs below 1%, and maintained consistent mass determination across the concentration and solution conditions examined. Controlled mixtures of antibody-derived species showed a linear molar-ratio-dependent ion response, supporting semi-quantitative estimation of relative species abundance under defined conditions. For heat-stressed trastuzumab, CDMS measurements were qualitatively consistent with SEC-MALS, mass photometry, and conventional native MS, while enabling direct mass determination of individual ions within highly heterogeneous high-molecular-weight (HMW) populations that exhibited extensive charge-state overlap and could not be reliably deconvoluted by conventional native MS. High-duty-cycle acquisition increased sampling of low-abundance, high-*m/z* ions and improved access to higher-order aggregate signals. Joint single-ion mass–charge analysis further differentiated statistical ion families within the broad HMW fraction and related major mass differences to independently obtained compositional information. These results support commercial Xevo^TM^ CDMS as a practical multidimensional platform for antibody aggregate characterization.

## 1. Introduction

Monoclonal antibodies (mAbs) constitute a major class of therapeutic biologics and are widely used in the treatment of cancer, autoimmune diseases, infectious diseases, and other complex disorders.^1–4^ However, aggregation, fragmentation, and other forms of molecular heterogeneity can arise during manufacturing, formulation, storage, and handling. Because these product-related variants may affect biological activity, pharmacokinetics, and immunogenicity, their accurate characterization is important for biopharmaceutical quality control and stability assessment.^5–9^ Higher-order aggregates are particularly challenging because increasing molecular mass, polydispersity, and compositional heterogeneity can obscure low-abundance species and reduce complex populations to ensemble-averaged descriptors.

Size-exclusion chromatography coupled with multi-angle light scattering (SEC-MALS) is widely used for characterizing antibody size variants and provides absolute molar-mass information for chromatographically separated populations.^10–13^ Sedimentation-velocity analytical ultracentrifugation (SV-AUC) provides an orthogonal, column-free approach for assessing aggregate distributions in solution ^14^ Mass photometry (MP) further enables rapid, label-free single-particle mass measurements with low sample consumption and has been applied to antibodies and heterogeneous immune complexes.^15^ Emerging mass-sensitive particle-tracking approaches additionally combine optical mass measurements with Brownian-motion-derived diffusion information, providing an orthogonal physical descriptor of individual particles ^16^ Native mass spectrometry (native MS) has already been proven successful on conventional MS detectors by providing complementary molecular mass and structural information via electrospray ionization under non-denaturing conditions.^17–19^ However, for highly heterogeneous, polydisperse, or megadalton-scale protein assemblies, extensive overlap of charge-state distributions can complicate charge-state assignment and conventional spectral deconvolution.^20, 21^ Thus, these approaches provide complementary views of aggregate populations, but highly heterogeneous HMW fractions can remain difficult to resolve at the constituent-population level.

Charge detection mass spectrometry (CDMS) addresses this analytical challenge by independently measuring the mass-to-charge ratio (m/z) and charge of individual ions, enabling direct mass determination without requiring resolved charge-state envelopes.^22, 23^ This single-ion approach has been applied to heterogeneous high-mass assemblies, including antibodies, immune complexes, and antibody-derived aggregates.^24–28^ Previous CDMS studies further showed that joint mass–charge information can help distinguish heterogeneous subpopulations,^29^ reveal conformational differences in megadalton antibody aggregates ^30^ and support relative-abundance measurements, including through

Direct Mass Technology.^31^ Nevertheless, detailed subpopulation-level characterization of highly heterogeneous higher-order therapeutic antibody aggregates remains less developed than that of lower-order oligomers, particularly dimers ^32^ Against this background, subpopulation-resolved interpretation of complex higher-order antibody aggregates remains comparatively underexplored,^33, 34^ particularly when multiple oligomeric states and chain-derived components coexist within a broad HMW distribution.

In this Technical Note, a ELIT-based CDMS platform was benchmarked for therapeutic mAb analysis and applied to heat-stressed trastuzumab. Orthogonal measurements provided compositional context, while joint single-ion mass–charge analysis with Gaussian mixture modeling was used to resolve heterogeneous HMW statistical ion families. Controlled mixtures with predefined molar ratios further evaluated the potential of CDMS for semi-quantitative assessment of relative abundances of species based on known molar ratios. Together, these results demonstrate the utility of commercial Xevo^TM^ CDMS for multidimensional characterization of heterogeneous antibody aggregates.

## 2. Experimental Section

### Materials

Recombinant anti-HER2 humanized monoclonal antibody (trastuzumab, mAb-1) and human-mouse chimeric anti-tumor necrosis factor-α monoclonal antibody (infliximab, mAb-2) were both manufactured and generously provided by Zhejiang Bioray Biopharmaceutical Co Ltd (Zhejiang, China) and WuXi Biologics (Jiangsu, China). The detailed composition of the original formulation is provided in Table S1. Bovine albumin (BSA, ≥99.9%) was purchased from Sigma-Aldrich (St. Louis, MO, USA). Enzyme-free water, Tris(hydroxymethyl)aminomethane (Tris), methanol, Sodium Dodecyl Sulfate (SDS), acetonitrile, trifluoroacetic acid (TFA) and ammonium acetate (5 M, LC-MS grade) were all purchased from ThermoFisher Scientific (Waltham, Massachusetts, USA). Micro Bio-Spin P-6 Gel Columns were purchased from Bio-Rad Laboratories. Inc. (Hercules, CA, USA).

### Sample pretreatment

First, the mAb-1 sample was dissolved in 20 mM histidine buffer supplemented with 8% (w/v) sucrose at pH 5.8, and subsequently subjected to thermal stress at 60 °C for one week. Semi-preparative SEC was utilized to collect fractions with distinct molecular weights. SEC was performed on an Agilent 1260 HPLC system equipped with a TOSOH TSKgel G3000SWXL column (7.8 mm × 300 mm, 5 μm; Tokyo, Japan). All SEC fractions were collected according to the predefined collection windows, followed by further buffer-exchange and concentration. Subsequently, all fractions were stored at −80°C for long-term storage prior to subsequent cross-validation and characterization by MP, native MS, and CDMS.

### SEC fractionation and SEC-MALS detection

The collected SEC fractions were further analyzed by SEC-MALS using a DAWN^TM^ MALS System connected to a 1260 Agilent HPLC. Each fraction was injected directly onto a TOSOH TSKgel G3000SWXL (5 μm, 7.8 mm * 30 cm) column for analysis. Data were acquired and processed with ASTRA^TM^ Software (Wyatt Technology, Santa Barbara, CA, USA). Absolute molecular masses were calculated from light scattering and RI signals according to Zimm’s equation. The weight-average molecular mass (Mw) and the relative mass fraction of HMW aggregates, monomer, and LMW species were determined for each SEC fraction.^13, 35^

### CDMS detection and method validation

All experiments were performed on a Waters Xevo™ CDMS mass spectrometer (Waters Corporation, Wilmslow, UK). Glass emitters for nanoelectrospray ionization (nanoESI) were fabricated from glass capillaries using a horizontal micropipette puller. The resulting emitter geometry was evaluated by cold-field-emission scanning electron microscopy, yielding a tip diameter of approximately 2 μm. Samples were loaded into the manual injection line at the front end of the mass spectrometer using a 25 μL gas-tight syringe (Hamilton Company, Reno, NV, USA).

All measurements were acquired in positive-ion mode. Analytes were ionized by nanoESI and transmitted sequentially through the StepWave™ ion guide, beam attenuator, segmented quadrupole, and ion-transfer lens assembly before entering the electrostatic linear ion trap (ELIT). The capillary voltage was set to 0.85–1.4 kV, the inlet temperature was maintained at 25 °C, and the peak-to-peak RF voltages were set to 250 V for the StepWave, 350 V for the ion guide, and 200 V for the segmented quadrupole. The ion trapping time was varied from 100 to 1200 ms, as appropriate. Prior to each acquisition, ion flux to the ELIT was optimized by adjusting the capillary voltage and beam attenuation to maintain predominantly single-ion trapping conditions; these settings were then kept constant during the acquisition.^36^

High duty cycle (HDC) acquisition mode ^36, 37^ was employed to enhance detection of low-abundance antibody aggregates across the *m/z* 8000–16000 range. During each ELIT trapping event, incoming ions were accumulated in the upstream segmented-quadrupole cooling region. Following completion of the trapping event, the accumulated ions were released toward the ELIT. Because ions of different *m/z* values exhibit different transit times, defined trapping-delay sequences enabled preferential trapping of ions within the target *m/z* range. This approach increased sampling efficiency for low-abundance, high-*m/z* ion populations without implying complete exclusion of ions outside the targeted range.

Analytical performance was benchmarked using representative attributes including mass and charge accuracy, repeatability, intermediate precision, practical concentration range, and robustness to selected solution conditions.

### Mass-Conditioned charge and Gaussian mixture analysis

Gaussian mixture modeling (GMM) with expectation–maximization was implemented in the Anaconda environment (Anaconda Software Distribution, Austin, TX, USA) using full-covariance Gaussian components ^29^ Two complementary analyses were conducted. First, the raw H1 single-ion mass–charge distribution was fitted with models containing 1–20 components. Although BIC favored an eight-component model, a 20-component fit was retained as a descriptive representation of local structure within the broad distribution. Components with fitted abundances >1% were displayed, yielding 17 major statistical ion families. Here, a statistical ion family refers to a group of ions assigned to the same Gaussian component on the basis of similar joint mass and charge characteristics. These components were not interpreted as discrete molecular species without independent compositional evidence.

Second, a mass-conditioned analysis was used to assess whether ion charge retained population-specific information beyond its expected dependence on molecular mass. Under native electrospray conditions, the average charge of compact proteins and protein complexes can be approximated by an empirical power-law relationship,*z=Cm^b,^*^38^ consistent with charged-residue-based scaling behavior.^39^ Because H1 contains highly heterogeneous antibody aggregates,*C* and *b* were fitted directly from the H1, H2 and H3 single-ion data rather than fixed to literature values.

For ions, logarithmic transformation gives:

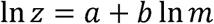

Where *a=*In*C*, and *m* and *z* denote the measured mass and charge, respectively. A mass-conditioned charge residual was then calculated as:

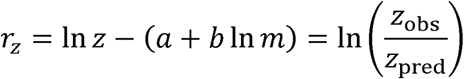

such that positive and negative values indicate charging above or below that expected for ions of comparable mass. GMM was subsequently performed in the (In *m, rz*)space after standardization of both variables. Full-covariance models containing 1–20 components were evaluated, with BIC used for formal model selection; a nine-component model was selected. Model stability was assessed across multiple initializations using the adjusted Rand index. The full nine-component model was retained for analysis, while the minor C5 component (0.74%) was omitted from selected visualizations for clarity. Components were interpreted as statistical ion families rather than discrete molecular species.

## 3. Results and Discussion

### Performance Benchmarking of Commercial Xevo^TM^ CDMS for Antibody Analysis

Because the fundamental mass and charge measurement performance of Xevo^TM^ CDMS has been established previously,^37^ baseline accuracy and precision were first confirmed before evaluating antibody-relevant operating conditions. Using β-galactosidase, a measured mass of 468.7 kDa differed by only 0.2% from the reference value of 468.76 kDa, while the charge measurement showed an RMS error below 0.2 e. BSA monomer and dimer measurements further showed repeatability and intermediate-precision RSDs below 1% for both mass and charge (Figure S1 and Tables S2–S5).

Practical performance for antibody analysis was subsequently evaluated across sample concentration and solution conditions. Concentrations of 0.005–0.01 mg/mL (33–67 nM) provided reproducible single-ion distributions within the fixed acquisition period, whereas 0.001 mg/mL yielded insufficient ion statistics within the acquisition period and 0.1 mg/mL required stronger control of ion flux (Figures 1A, S2 and S3). Variation in pH and ammonium acetate concentration affected charge distributions and ion flux more strongly than the measured mass (Figures S4-S6), and reproducible mass measurements were retained under the selected native-like conditions. Based on these results, 200 mM ammonium acetate was used for subsequent analyses.

**Figure 1.**
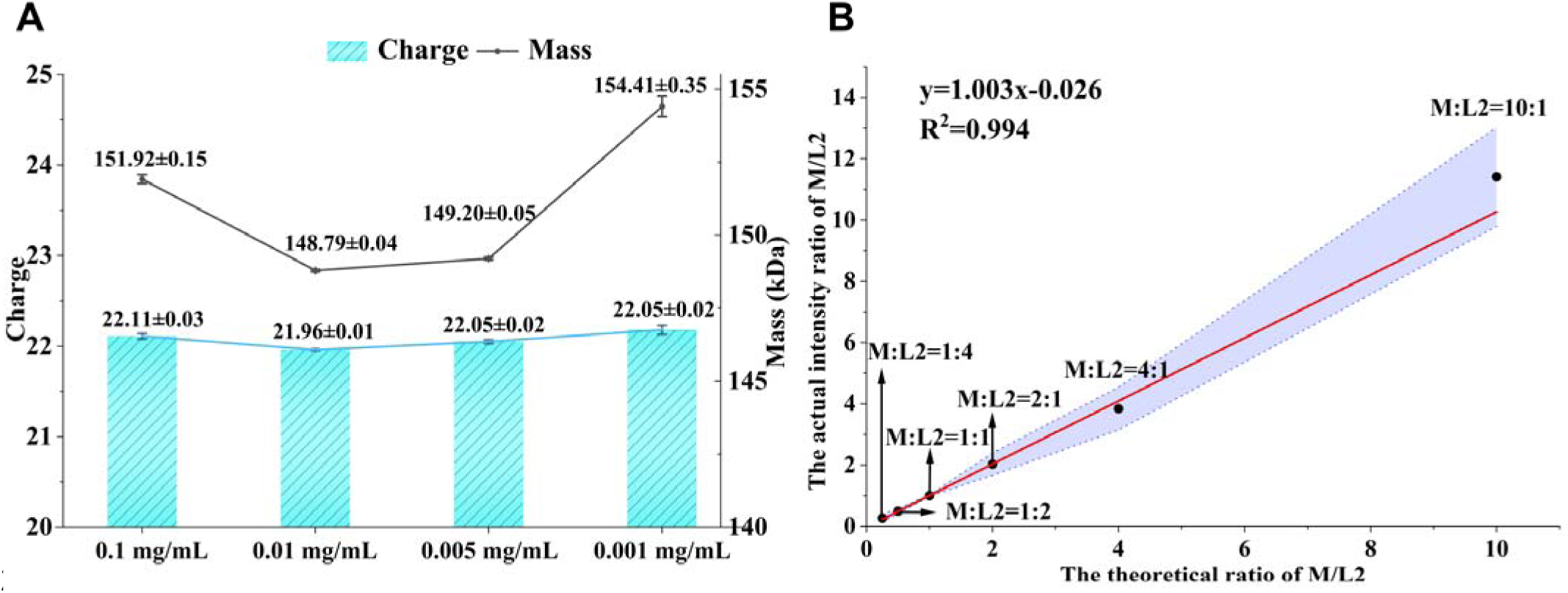
Charge and mass distribution profiles of the antibody samples. Bar charts represent charge state distributions, while scatter plots represent mass distributions. (A) mAb-1 at varying mass concentrations. All datasets were derived from Gaussian fitting of raw mass and charge measurements. (B) Linear fit analysis of CDMS peak area-weighted intensities across different molar ratios. The x-axis represents the nominal molar fraction calculated from the prepared Main Peak/LMW2 (M/L2) mixtures, whereas the y-axis represents the corresponding CDMS-derived relative ion abundance. The blue area in the graph indicates the error bars. The data points denote the measurements acquired at various molar ratios, specifically 10:1, 4:1, 2:1, 1:1, 1:2, and 1:4 (n=3).

Recent studies have demonstrated the quantitative potential of single-ion CDMS for heterogeneous protein systems ^31^ In the present study, the applicability of this principle to antibody-derived species was further evaluated by examining whether CDMS-derived relative ion abundances could reproduce predefined molar ratios over a controlled mixing range. Controlled mixtures of SEC-separated LMW2 (L2) and main peak (M) fractions were therefore prepared at predefined molar ratios for experimental evaluation. These two fractions were selected as model analytes owing to their high structural stability, negligible non-covalent depolymerization behavior, and high chromatographic purity.

Antibody mixture samples were prepared at predefined molar ratios of M to L2, spanning ratios of 1:1, 2:1, 1:2, 1:4, 4:1, and 10:1. All mixtures were analyzed by CDMS, and the resulting data were subjected to purity correction and weighted normalization, as presented in Figure 1B. A coefficient of determination (R²) greater than 0.99 and slope 1.003 were achieved, with minor deviations observed only at the extreme 10:1 ratio. These results support a molar-ratio-dependent response of CDMS-derived abundance under the tested conditions. This behavior differs from the mass-weighted response typically obtained from concentration-based measurements such as SEC-UV ^40^ or

SEC-MALS, which yield only average molecular weight information and lacks direct particle counting capability. Therefore, single-ion CDMS has the potential for molar-population-based semi-quantitation for heterogeneous macromolecular species. However, it is well known that ionization and transmission efficiency can vary based on analyte physiochemical properties and instrument tuning in MS based methods. Analytical performance in practical laboratory workflows may be influenced by a broad spectrum of confounding and matrix-related interfering factors ^41^ Therefore, appropriate calibration strategies should be employed for reliable quantitation.

### Enhanced Dynamic Range of CDMS Resolved Heterogeneous Higher-Order Aggregates in heat-stressed mAb-1

A thermal stressed mAb-1 was characterized by SEC, MP, conventional native MS, and CDMS (Figures 2A-D and S7). All methods showed noticeable degradation into both lower and higher MW species than the original mAb mononer at 150 kDa. SEC, performed before buffer exchange, resolved three HMW peaks, the main monomer peak, and two LMW fragment peaks, with HMW2 corresponding approximately to the antibody dimer. The other methods also identified signals from dimers and lower molecular weights. Despite the consistent qualitative detection of the major species in the sample, the quantitative information varied significantly among the different methods. This is at least partly due to the substantially different sample concentrations and preparation conditions. SEC required the highest concentration of 67 μM. In contrast, CDMS, MP, and native MS were performed after buffer exchange and at substantially lower analytical concentrations (33 nM, 5 nM, and 1000 nM, respectively). Dilution reduces the absolute concentration of low-abundance aggregate species and may also shift the equilibrium of reversible non-covalent assemblies toward dissociation. This effect is particularly relevant to highly diluted measurements and has been recognized as an important consideration in single-particle analyses of non-covalent protein complexes.^42^ Consequently, the distributions observed after buffer exchange should be interpreted quantitatively as those retained under the respective measurement conditions rather than as direct representations of the original formulation. Nonetheless, MP and MS based methods have higher resolving power and can provide critical qualitative information to explain the low-abundance, high-mass species seen in SEC-MALS.

**Figure 2.**
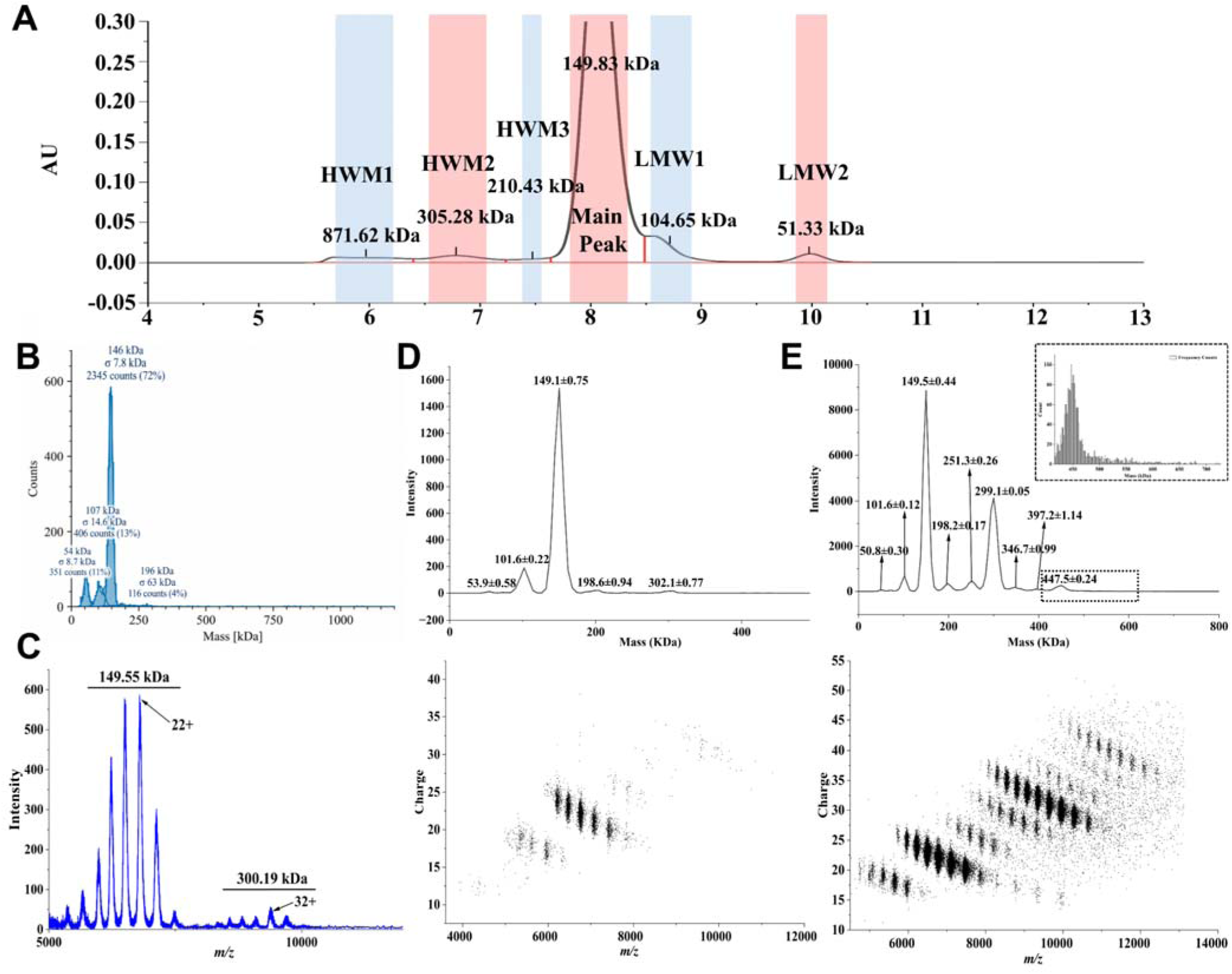
Multi-instrument characterization of full samples after thermal stress. (A)SEC chromatogram and the corresponding collected fraction samples. Peaks are assigned from left to right as high molecular weight 1 (HMW1), HMW2, HMW3, main peak, low molecular weight 1 (LMW1), and LMW2, with their respective retention times labeled adjacent to each peak. (B)MP; and (C) native MS. The low *m/z* were filtered out in the method for qualitative analysis and improved sensitivity on high *m/z*. (D) CDMS under minimally activating conditions; (E) CDMS under HDC mode. The black dotted box in the image represents a zoom-in of the selected area. Top: Mass spectrum; Bottom: *m/z* vs. charge 2D plot. The mass values in the mass spectrum were obtained after Gaussian fitting. All results without SEC were measured after the buffer was exchanged.

To enhance sampling efficiency for low-abundance ions in the high-m/z region, the high duty cycle (HDC) mode of CDMS (shown in Figure 2C) was applied to improve ion utilization efficiency (duty cycle).^37^ Compared with the standard acquisition, HDC improved visualization of higher-mass ion populations and resolved discrete oligomeric bands through the mAb-1 trimer, effectively matching the expected MW in HMW1 in the SEC-MALS data. Similar m/z-selective accumulation strategies have been used on trapping-based platforms, whereas conventional TOF analyzers generally require additional upstream gating or accumulation hardware for comparable duty-cycle enhancement ^43^ This highlights the flexibility of Xevo^TM^ CDMS for extending access to higher-order antibody aggregates without prior fractionation. For MP, expansion of dynamic range using such tuning is not readily accessible. This demonstrated the flexibility of the CDMS system to rapidly assess higher MW mAb aggregates without the need of pre-fractionation. Based on this analysis, it is estimated that ≥3 orders of magnitude improvement in dynamic range can be achieved simply by utilising the HDC functionality. However, the distribution above the trimer (∼500 kDa) remained unresolved and are likely highly heterogeneous, which may correspond to HMW1 in the SEC-MALS. To further improve the dynamic range, SEC fractionations were therefore collected and analyzed separately.

### Complementary Data identified H1 as a highly heterogeneous HMW fraction

As illustrated in Figure 2A, thermal stressed samples were subjected to SEC separation, and six discrete elution fractions were collected for subsequent analytical experiments. For concise reference in the following discussion, these fractions are designated with simplified abbreviations: high molecular weight fractions 1–3 (HMW1–3) are referred to as H1–H3, the main peak as M, and low molecular weight fractions 1–2 (LMW1–2) as L1–L2.

The collected SEC fractions were further characterized by SEC-MALS (Table 1 and Figures S9–S13), MP (Figure S14), native MS on TOF (Figures S15 and S16), and CDMS (Figure S17). Again, all techniques showed good consistency qualitatively on the species present in each fraction. However, as MP and MS-based methods required samples at significantly lower concentration than SEC-MALS, all fractions showed low MW species which may have dissociated from higher-order complexes (Figures S14 and S15). In addition, low-mass MP signals can be susceptible to background-related artifacts, which may complicate interpretation of weak fragment populations ^44^ Herein, we only focus on using these data to assign the MW of the SEC fractions. The H1 fraction was not well resolved by any of the techniques. In SEC-MALS, H1 showed an average molecular weight of 871.62 kDa and a higher dispersity than the remaining fractions, indicating that H1 contains a substantially broader molecular-weight distribution rather than a single well-defined oligomeric species, consistent with broad mass distribution seen by MP and MS-based methods. SEC-MALS is well suited for determining weight-average molecular masses and assessing the overall dispersity of chromatographically separated populations. However, when multiple higher-order species co-elute within a broad SEC fraction, the measured molecular weight represents the collective distribution and does not directly identify the molecular masses or stoichiometries of the individual constituent species.^45^

**Table 1.** SEC fraction samples and their key parameters. Purity values are those prior to buffer exchange, which are the values from the original formulation.

| Fraction Name | Retention Time (min) | Average Molecular Weight (kDa) | Polydispersity (Mw/Mn) | Purity (before buffer exchange, %) | Area (%) |
| --- | --- | --- | --- | --- | --- |
| H1 | 5.70 | 871.62 | 1.07 | 96.1 | 2.81 |
| H2 | 6.89 | 305.28 | 1.00 | 92.4 | 2.94 |
| H3 | 7.58 | 210.43 | 1.00 | 91.2 | 0.76 |
| M | 8.16 | 149.83 | 1.00 | 99.8 | 86.19 |
| L1 | 8.71 | 104.65 | 1.00 | 45.1 | 5.17 |
| L2 | 10.09 | 51.33 | 1.00 | 93.9 | 1.99 |

Native MS provided well-resolved mass information for the lower-complexity fractions, including the LMW species at approximately 48.21 and 100.17 kDa, but the extensive charge-state overlap of H1 precluded reliable deconvolution (Figure S15). In contrast, CDMS achieved direct mass and charge measurements across the SEC fractions without requiring resolved charge-state envelopes (Figure S17). Thus, the orthogonal measurements consistently identified H1 as the most heterogeneous fraction, while CDMS preserved single-ion information within a mass range that remained poorly resolved by conventional deconvolution.

To provide compositional context for the HMW populations, H3 and H1 were further examined using native MS fragmentation, denaturing RP LC-MS, and nrCE-SDS. CID and SID of the lower-complexity H3 fraction produced fragments near 23.8 and 47.2 kDa, and RP LC-MS identified light chain (LC)-derived species, LC dimers, light chain – heavy chain ([LC][HC]), intact mAb, and related components (Figures S18–S20A and S21; Table S6). H1 was less amenable to CID/SID, but RP LC-MS similarly revealed LC, LC dimer, [LC][HC], and intact mAb-derived components, while nrCE-SDS was consistent with contributions from both chain-derived species and noncovalent association (Figures S20B, S22 and S23; Table S7).^46^ Notably, [LC][HC] was detected in H1 and H3, indicating that the HMW populations contain building blocks beyond intact-mAb and LC-dimer units. These measurements define the principal molecular building blocks contributing to the HMW fractions but, because they involve dissociation or denaturation, do not preserve their correspondence to the intact higher-order assemblies. This distinction motivated subsequent analysis of the H1 population using the joint single-ion mass and charge information retained by CDMS.

### Joint Single-Ion Mass–Charge Analysis Resolves Heterogeneous Higher-Order Aggregate Populations

Based on the lower-MW fractions and reported antibody fragmentation behavior, a plausible degradation–aggregation scheme involving hinge-region fragmentation and subsequent incorporation of LC-, LC-dimer-, and [LC][HC]-related components into heterogeneous higher-order assemblies is proposed in Figure 3. Such compositional complexity provides a plausible basis for the unusually broad H1 distribution but cannot be reconstructed directly from measurements performed after dissociation or denaturation. CDMS therefore provides a complementary advantage by retaining the mass and charge of each intact detected ion.

**Figure 3.**
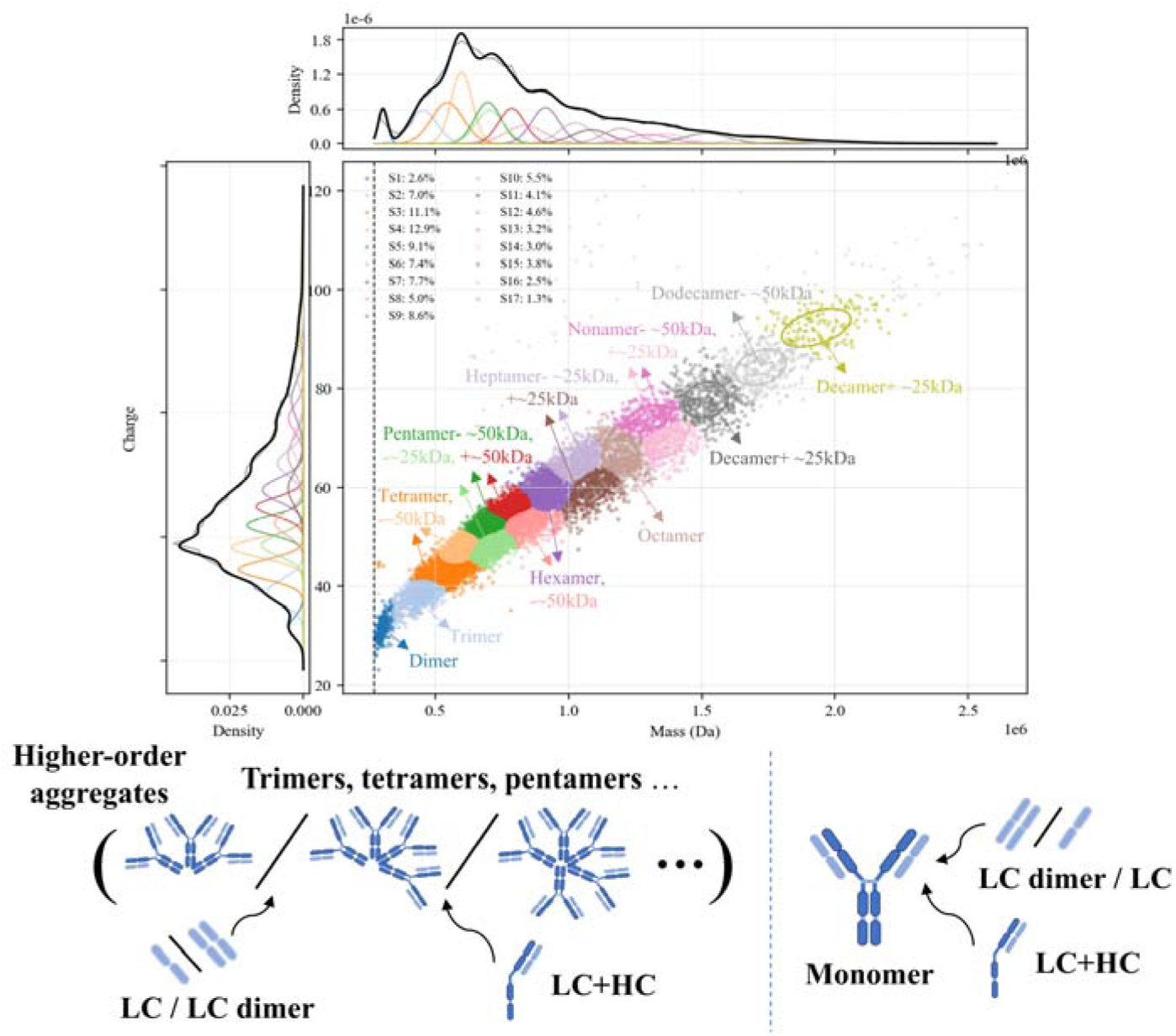
Top Two-dimensional mass versus charge plot showing ion families obtained from Gaussian fits to the experimental data for the CDMS measurement of H1 fraction. See main text for details. This model finds evidence for seventeen ion families with abundances greater than 1%. Each point shows the mass and charge measured for a single particle and is assigned to an ion family (indicated by color). Subfamily assignment is based on the highest probability of each particle belonging to a specific subfamily. Visually, this leads to boundaries that are artificially strict as, in reality, the ion families overlap. The top and left side traces show the integrated raw data for the mass and charge dimensions respectively and corresponding sums of the Gaussian curves as black lines for these dimensions. The determined fits for each ion family are also delineated using the same color scheme. The percentage of each ion family is also indicated. The bottom schematic summarizes experimentally supported compositional building blocks and a proposed relationship to the HMW population, it is not intended to assign individual GMM components to unique molecular structures.

The H1 fraction exhibited extensive heterogeneity in both dimensions, and projection onto either mass or charge alone produced strongly overlapping distributions (Figure 3). Two-dimensional Gaussian mixture modeling (Figure S24) was therefore used to describe the population structure in the joint mass–charge space rather than to assign individual molecular species. Seventeen major fitted ion families with relative abundances above 1% were resolved, spanning approximately 302 kDa to 1.95 MDa with charge centroids of approximately 32–93 e (Table S8). This behavior is consistent with previous CDMS studies showing that jointly measured mass and charge can provide greater statistical separation of heterogeneous macromolecular populations than either one-dimensional projection alone.^47^ Importantly, neighboring H1 populations frequently differed in both mass and charge, indicating that the charge dimension contributes information beyond a simple mass spectrum.

The mass centroids also provided a link to the complementary compositional measurements. Recurrent offsets of approximately 25 and 50 kDa were observed among neighboring populations, consistent with contributions from LC- and LC-dimer-related species identified by RP LC-MS and fragmentation analyses. RP LC-MS and nrCE-SDS additionally supported the presence of a [LC][HC] species. A distinct ∼75 kDa spacing was not resolved as an independent H1 GMM population. However, this does not exclude its contribution to the intact aggregates. Several major H1 populations have mass widths of approximately 30–55 kDa (Table S8), comparable to or exceeding the ∼25 kDa difference between 50 and 75 kDa compositional increments. Closely related aggregate variants may therefore be represented within the same broad Gaussian population. Accordingly, the observed mass offsets are best regarded as compositional signatures rather than unique stoichiometric assignments, and the fitted components are interpreted as compositionally related ion families.

Although the two-dimensional analysis demonstrates that charge contributes to population separation, absolute charge is intrinsically coupled to molecular mass: larger protein assemblies generally carry more charge under native electrospray conditions ^48^ Consequently, part of the apparent separation along the charge axis in Figure 3 may simply reflect this mass-dependent charging trend rather than population-specific charging behavior. To determine whether charge retains discriminatory information after this expected dependence is removed, a second analysis was performed using mass-conditioned charge.

The global mass–charge relationship was modeled in logarithmic space, and the deviation of each ion from its mass-predicted charge was expressed as a mass-conditioned charge residual. Residual-GMM analysis revealed partially overlapping ion families with distinct mass-normalized charging behavior (Figure S25 and Table S9). Thus, even among ions of comparable molecular mass, measurable differences in charging remained after removal of the dominant mass dependence. For example, C3 and C4 were centered at approximately 612 and 673 kDa, respectively, but showed residual-charge centroids of +0.055 and −0.069, corresponding to observed charge levels approximately 5.7% above and 6.7% below those predicted from the global relationship. These differences suggest that factors beyond molecular mass—potentially including conformation, surface accessibility, or composition—contribute to charging, although residual charge alone does not provide a direct structural assignment.

Together, the two analyses provide complementary levels of information. The joint mass–charge GMM describes the intact H1 population and relates major mass differences to independently identified compositional features, whereas the mass-conditioned analysis tests whether charge contains additional population information beyond its expected dependence on molecular size. This combination extends CDMS beyond direct mass measurement by providing a multidimensional, single-ion view of higher-order antibody aggregate heterogeneity that is difficult to obtain from ensemble-averaged mass measurements or conventional charge-state deconvolution.

## 4. Conclusions

This study establishes the practical performance of commercial Xevo^TM^ CDMS for therapeutic antibody aggregate analysis. The platform provided reproducible mass and charge measurements and maintained consistent mass determination across the concentration and solution conditions examined. For heat-stressed trastuzumab, the major populations observed by CDMS were broadly consistent with SEC-MALS, MP, and native MS, while direct single-ion measurement avoided the requirement for resolved charge-state envelopes. Importantly, high-duty-cycle acquisition increased the sampling efficiency of low-abundance high-*m/z* ions and extended the effective accessible range toward higher-order aggregates. This tunable acquisition capability is particularly useful for antibody samples in which abundant monomer and low-abundance HMW species coexist over a wide mass and abundance range.

The highly heterogeneous H1 fraction further illustrates the additional information available from CDMS beyond direct mass determination. Joint mass–charge analysis provided population-level differentiation within a broad HMW distribution, while complementary compositional measurements supported contributions from intact mAb and chain-derived species. These statistical populations should not be interpreted as unique molecular assignments, but they demonstrate how the single-ion mass and charge dimensions can complement established aggregate-characterization methods. Controlled mixtures also indicated that ion abundance can approximately reflect prepared molar ratios under defined conditions, although broader quantitative application will require appropriate calibration and control of ionization and transmission effects. Future coupling of CDMS with online, native-compatible separation (i.e., SEC ^28^, field-flow fractionation ^49^) could further improve the analysis of complex antibody mixtures by combining chromatographic resolution with direct single-ion detection, while reducing the need for offline fraction collection and minimizing perturbation of labile aggregate populations.

## Acknowledgments

Funding was generously provided by the Huadong Medicine Joint Funds of the Zhejiang Provincial Natural Science Foundation of China (Grant No. LHDMZ24H300003), National Natural Science Foundation of China (Grant No. 82674934), Zhejiang Provincial Science and Technology Project for Marine Industry (Grant No. 2026HYC10005). This work was also supported by the Instrument Innovation Program of Institute of Fundamental and Transdisciplinary Research, Zhejiang University and the Protein Characterization and Crystallography Facility of Westlake University. We thank Ms. Haihong Hu for laboratory management at Zhejiang University and Mr. Qi Shen from Waters Co., Ltd (Shanghai, China).

## Declaration of generative and Al-assisted technologies in the manuscript preparation process

During the preparation of this work the authors used Chatgpt-5.6 in order to polish language. After using this tool, the authors reviewed and edited the content as needed and took full responsibility for the content of the published article.

## Authorship contribution statement

**Shang-Yin Wu:** Writing - original draft, Methodology, Investigation, Data curation. **Xin-Miao Guo:** Investigation, Methodology. **Jia-Yi Lv:** Investigation, Methodology. **Ying-Zi Li:** Investigation, Methodology. **Lei Gu:** Investigation, Methodology. **Zachary Tian:** Investigation, Methodology. **Yong-Sheng Xiao:** Investigation, Methodology, conceptualization, methodological design, project administration, funding acquisition. **Emily Christofi:** Writing-review & editing, Investigation, Methodology. **Anisha Haris:** Writing-review & editing, Investigation, Methodology. **Haoyun Lu:** Investigation, Methodology. **Rui Wang:** Investigation, Methodology. **Huan-Fang Xie:** Investigation, Methodology. **Hongwen Li:** Investigation, Methodology. **Yuanjiang Pan:** provision of resources, funding acquisition. **Mowei Zhou:** Writing-review & editing, project administration, conceptualization, methodological design, provision of resources, and funding acquisition. **Wei-Jie Fang:** Writing-review & editing, conceptualization, methodological design, provision of resources, project administration and funding acquisition. All authors have given approval to the final version of the manuscript.

