## Supporting Information for "Benchmarking the Performance of a Commercial Electrostatic Linear Ion Trap (ELIT) Charge Detection Mass Spectrometer For Higher-Order Antibody Aggregate Analysis"

^5^Waters Co., Ltd, Wilmslow, SK94AX, UK

### **Supplementary experimental methods:**

***MP detection***

All MP measurements were carried out in Refeyn Sample Well Cassettes (11mm × 15mm, Oxford, UK) on microscopy slides that had been cleaned by consecutive sonication in water, isopropanol, and water. Protein solutions (20 μl) were added to the gaskets containing 4 μl buffer and images were acquired for 60 s at 1 kHz (original), 100 Hz (integrated). All pre-stress samples (following buffer exchange), post-stress intact bulk samples, and collected SEC fraction samples were separately diluted to 100 nM, yielding a final measurement concentration of approximately 5 nM after 1:20 dilution. The molecular weight and percentage composition of each component were subsequently determined via Refeyn Two MP mass photometer (Oxford, UK).

***Denatured reversed-phase LC-MS***

The molecular mass analysis was performed on Agilent/1290 Infinity II system with an Agilent/PLRP-S column coupled with an Agilent/6230 TOF MS system. Collected data were analyzed by the PMI-Byos software and the measured molecular masses were compared with the theoretical molecular masses to confirm consistency.

***nrCE-SDS***

The SEC fractions HMW1 and HMW3 were analyzed by non-reduced capillary electrophoresis-sodium dodecyl sulfate (nrCE-SDS) to determine their apparent molecular weights and to assess whether the HMW species were covalent or non-covalent. Non-reduced CE-SDS is performed using a PA800 Plus CE system (Brea, CA, USA) with PDA (photo-diode array) detector. Fractions were concentrated to 0.5–1.0 mg/mL by ultrafiltration if necessary, then mixed with CE-SDS sample buffer (0.1 M Tris-HCl, 1% SDS, pH 9.0) to a final protein concentration of 0.1–1.0 mg/mL. To prevent free sulfhydryl-mediated disulfide scrambling during sample preparation, iodoacetamide was added to a final concentration of 10 mM and the mixture was incubated at 70 °C for 5 min or at 25 °C for 15 min in the dark. Samples were injected electrokinetically at 5 kV for 10 s, and separation was performed at a constant voltage of 15 kV with reversed polarity at 25 °C. The relative abundance of HMW species, monomer, and fragments was determined by corrected peak area normalization.

***Native MS detection***

Native MS experiments were performed using a Waters Synapt^TM^ XS System (Waters Corporation, Wilmslow, UK), modified with SID after the mass-selective quadrupole ^1^. Data were acquired in resolution mode in both positive and negative ion modes. Protein solutions were infused using a static nanoelectrospray in positive ion mode through a platinum wire at ∼0.9 kV and 30°C, employing in-house pulled borosilicate glass capillaries (ID 0.78 mm, Item No. BF100-78-10, Sutter Instruments, USA) using a shutter instrument pipet puller P-1000 (Novato, CA, USA). The cone voltage was set at 80 V, with a source offset of 4 V. For MS/MS experiments, manual fixed quadrupole profile was used to filter out lower *m/z* species, and dissociation was achieved either by CID using trap collision energy (Trap CE) or by SID. SID collision energy was controlled via both the trap collision energy and the collector/stopper voltage ^2^.

***Data analysis***

The resulting histograms of MP were fitted to Gaussian distributions using Refeyn Ltd. DiscoverMP (Oxford, UK) to extract the peak contrast and relative amount of each peak. Data analysis of native MS was conducted using MassLynx^TM^ Version 4.2 Software and UniDec 6.0.4 ^3^. The Gaussian fit of the CDMS mass distribution was performed in Origin 2026 (OriginLab, Massachusetts, USA) to determine the center-of-mass mass. The reported error corresponds to the standard deviation (σ) obtained from the Gaussian fit, reflecting the observed ion population width, which includes factors such as adduct heterogeneity, molecular heterogeneity, and uncertainties in charge measurements. Signal processing and visualization were performed using waters_connect^TM^ CDMS Toolkit Software.

### **Supplementary experimental results**:

***Accuracy of charge, mass, and mass-to-charge ratio***

This study employed β-galactosidase as a reference analyte to independently verify the accuracy of mass and charge assignments, thereby comprehensively validating the overall *m/z* measurement performance of the CDMS system. The experimentally determined mass of 468.7 kDa exhibited a relative deviation of 0.2% from the calibrated reference mass of 468.76 kDa. For charge state measurements, a root mean square (RMS) error of less than 0.2 e was achieved, with a mean deviation of 0.166 e. Collectively, these findings confirm that this CDMS platform delivers excellent analytical accuracy across mass, charge, and *m/z* determinations.

***Precision (including repeatability and intermediate precision)***

Given that BSA spontaneously forms aggregates during production, it was selected as the model protein for precision validation, enabling concurrent verification of mass and charge measurements for both monomeric and aggregated species. As illustrated in Figure S1, six replicate measurements yielded repeatability masses of 68.49 ± 0.15 kDa for BSA monomers and 136.30 ± 0.64 kDa for BSA dimers. Intermediate precision, evaluated by different operators on independent days, gave corresponding masses of 68.47 ± 0.11 kDa and 136.50 ± 0.83 kDa, respectively. Charge state measurements were consistent with the mass results, with relative standard deviation (RSD) values below 1.0% across all precision assays; complete datasets are provided in Tables S1–S4. Notably, all deviations in both repeatability and intermediate precision assessments were below 1.0%, confirming the excellent precision performance of the native CDMS platform.

***Influence of Solution Conditions on CDMS Response***

To further evaluate the analytical robustness and practical utility of CDMS, detection performance was systematically assessed across buffers with varying pH values and ammonium acetate concentrations. As shown in Figure S4A, initial validation experiments were conducted using mAb-1. Given the high inherent stability of mAb-1, which confers relative insensitivity to pH fluctuations. These results demonstrate that changes in sample and buffer conditions influenced the observed charge distributions and ion flux, while reproducible mass measurements were retained under the selected native-like conditions. To more rigorously challenge the analytical capabilities of CDMS, mAb-2 - a model antibody with moderately lower structural stability - was subsequently selected for further evaluation.

As presented in Figures S4 and S5, the optimal pH for mAb-2 was determined to be approximately 7. Under acidic conditions, acid-induced protein denaturation and fragmentation resulted in pronounced peak broadening and the emergence of ~100 kDa fragment species (Figure S5A). Counter to the expected shift toward higher charge states under acidic conditions, the charge distribution instead shifted toward lower charge values (Figure S5)^5, 6^. Meanwhile, compared with alkaline conditions, denatured protein ions are more readily enriched at the gas–liquid interface of electrospray droplets, yielding substantially higher ionization efficiency than natively folded protein and a marked increase in ion transmission efficiency^7^. This effect is detrimental to quantitative CDMS detection. Collectively, these findings support the recommendation that CDMS analysis be performed at pH values within ±1 unit of the protein’s optimal pH, with a slight preference for mildly alkaline conditions.

Results for varying salt concentrations are presented in Figure S3C and S5. With the exception of the 5 M condition, both mass and charge measurements yielded reliable results with no anomalous data. However, a marked shift in charge state distribution toward lower charge values (the charge reduction effect) was observed as salt concentration increased substantially ^8^. This phenomenon is likely attributable to ammonium ions competing for protein charge sites via gas-phase proton transfer reactions during ESI droplet evaporation, which reduces the average charge of the resulting protein ions. On this basis, 200 mM ammonium acetate was selected as the standard buffer background for buffer exchange in all subsequent experiments.

Taken together, these results demonstrate that while the spectral profiles of monomeric and aggregated protein species may undergo substantial alterations under acidic, alkaline, and high-salt stress conditions, CDMS maintains consistently stable and reliable detection performance across all tested conditions. These data further confirm the suitability of CDMS for analyzing stress-induced protein aggregation and evaluating protein stability.

***MP Reveals partial dissociation of HMW aggregates after dilution***

MP was used to characterize the six SEC fractions after buffer exchange (Figure 4) ^9^. The H1–H3 fractions showed partial loss of higher-order species, consistent with dissociation of non-covalent aggregates during dilution and buffer exchange. Because MP is typically performed at low nanomolar concentrations, reversible assemblies may be influenced by concentration-dependent shifts in association equilibria. In addition, the ~48 kDa L2 fragment lies near the lower-mass regime where MP detection becomes increasingly sensitive to background and experimental conditions ^10^. More importantly for the HMW fractions, MP provides a one-dimensional mass distribution and therefore cannot exploit charge as an additional descriptor when multiple heterogeneous aggregate populations overlap in mass. Thus, although MP provides valuable single-molecule information on mass distributions and aggregate dissociation, it does not resolve the compositional complexity of the H1 fraction. This limitation motivates subsequent analysis using CDMS, in which mass and charge are measured simultaneously for each detected ion.

***Native MS resolves lower-complexity species but not the heterogeneous H1 fraction***

To better understand the composition of the HMW1, we examined all the available data. In general, the stressed samples contain species combined from ~50 kDa and ~150 kDa. Native MS provided the most accurate mass information for fractions in which well-resolved charge-state distributions were retained (Figure 5 and Figure S13). The LMW fractions were dominated by species of 48.21 and 100.17 kDa. The measured mass of L2 was 48,210 Da. Based on the trastuzumab sequence, a species comprising the complete light chain and the N-terminal heavy-chain segment extending to P233 has a calculated mass of approximately 48,215 Da when the retained disulfide linkages and heavy-chain N-terminal pyroglutamate formation are considered. The close agreement supports assignment of L2 as a hinge-containing Fab-like fragment. This is consistent with previous reports ^11^. Notably, this species extends further into the hinge region than a conventional papain-generated Fab and contains hinge cysteines that participate in inter-heavy-chain disulfide linkages in intact IgG1. Therefore, its exact disulfide state cannot be established from intact mass alone. The major L1 species was observed at approximately 100,168 Da, consistent with an Fc–Fab-like complementary fragment generated by loss of one Fab-like arm. The complementary masses of the L1 and L2 species further support a common hinge-region fragmentation process, although the precise cleavage site and disulfide rearrangement cannot be assigned solely from intact native MS measurements. Together, these data indicate that thermal stress generates complementary Fab-like and Fc–Fab-like LMW species from mAb-1 ^12^. Converting the SEC data into molar ratios can also provide indirect confirmation of this.

### **Supplementary Figures:**

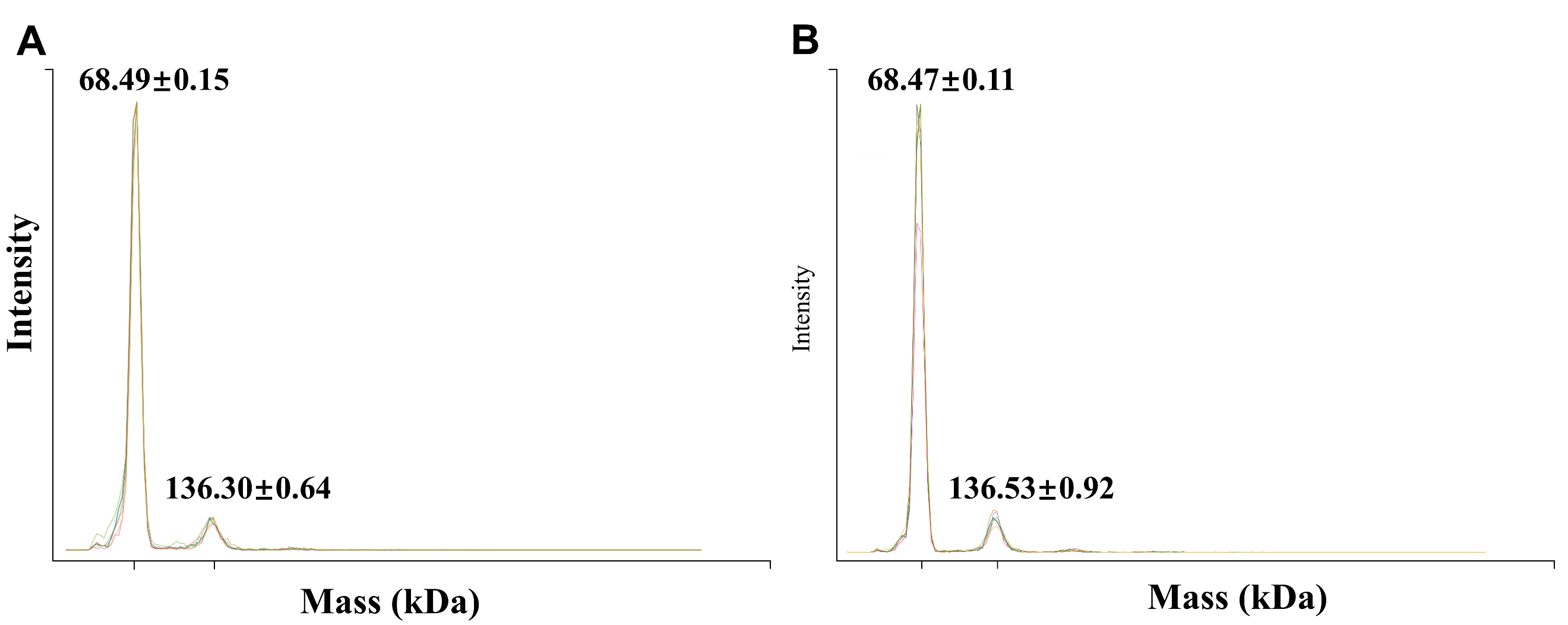

Figure S1. (A) Mass spectra of BSA from the repeatability study; (B) Mass spectra of BSA from the intermediate precision study. Both repeatability and intermediate precision were measured six times (the numbers in the figure represent the average mass obtained from the repeated measurements).

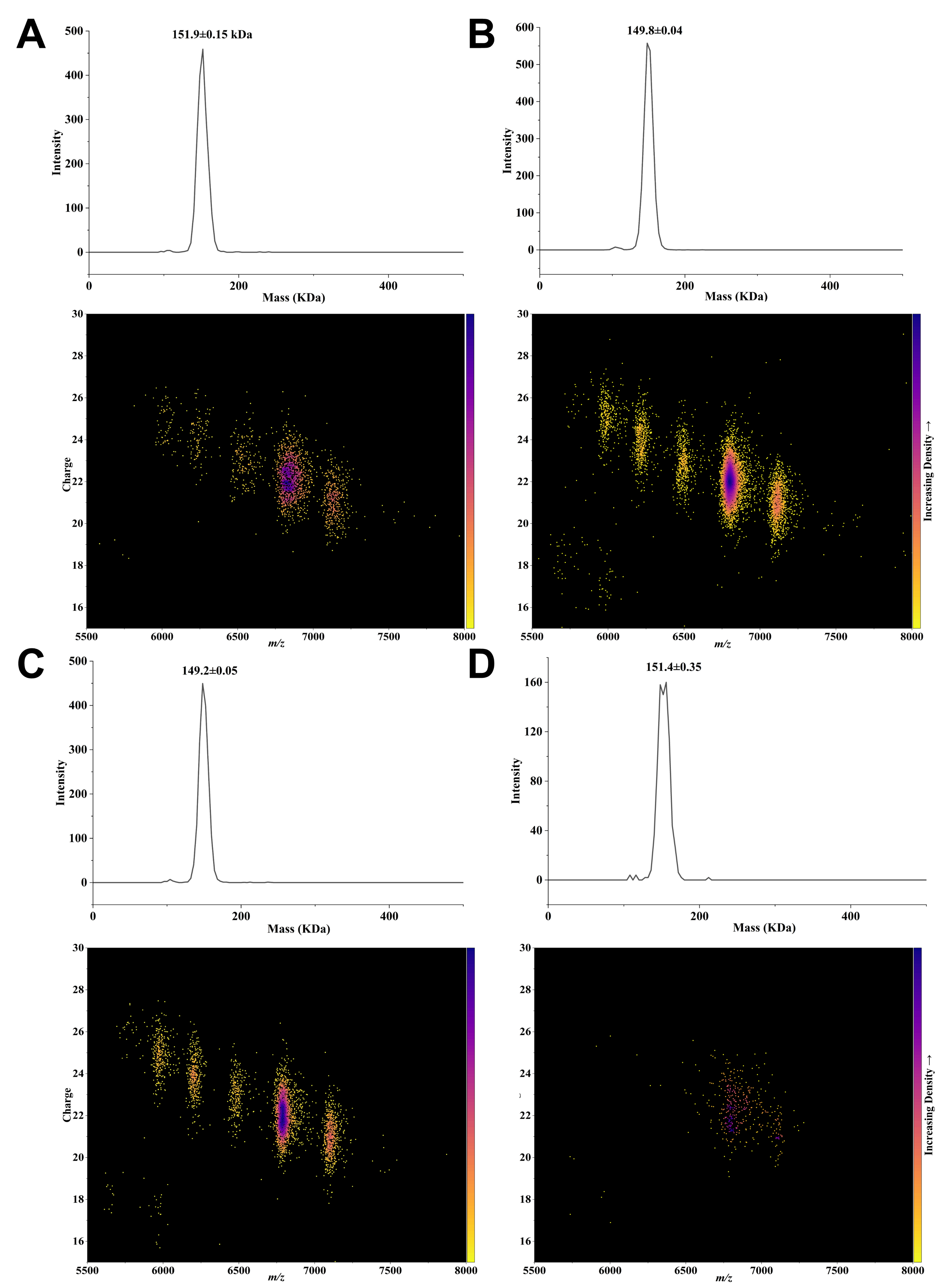

Figure S2. Native charge detection mass spectrometry of mAb-1 at different concentrations. (A) 0.1 mg/mL, (B) 0.01 mg/mL, (C) 0.005 mg/mL, and (D) 0.001 mg/mL. For each concentration, Top: Mass spectrum; Bottom: *m/z* vs. charge 2D point density plot. The mass values in the mass spectrum were obtained after Gaussian fitting.

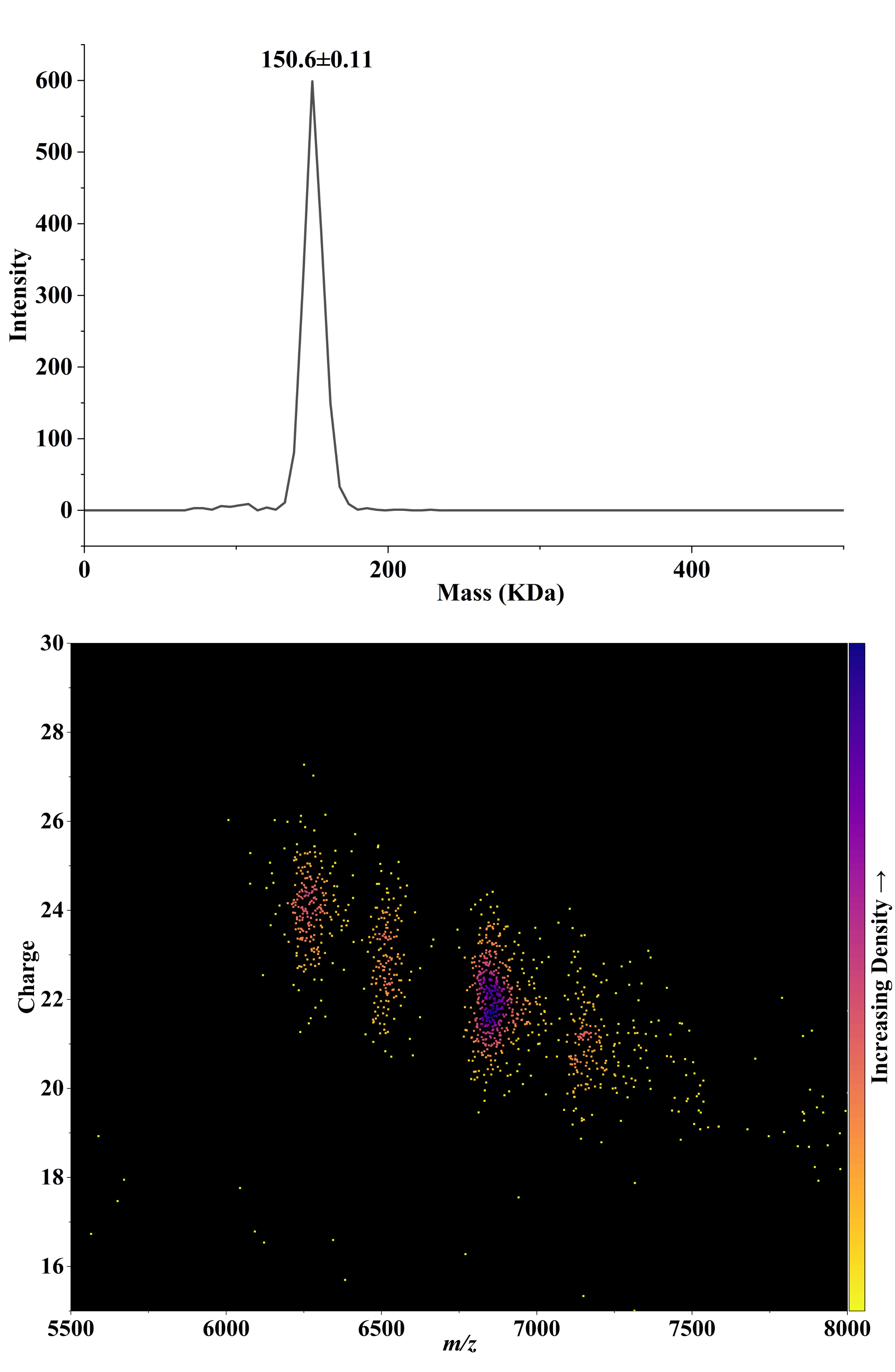

Figure S3. Native charge detection mass spectrometry of mAb-2 at 0.005 mg/mL. Top: Mass spectrum; Bottom: *m/z* vs. charge 2D point density plot. The mass values in the mass spectrum were obtained after Gaussian fitting.

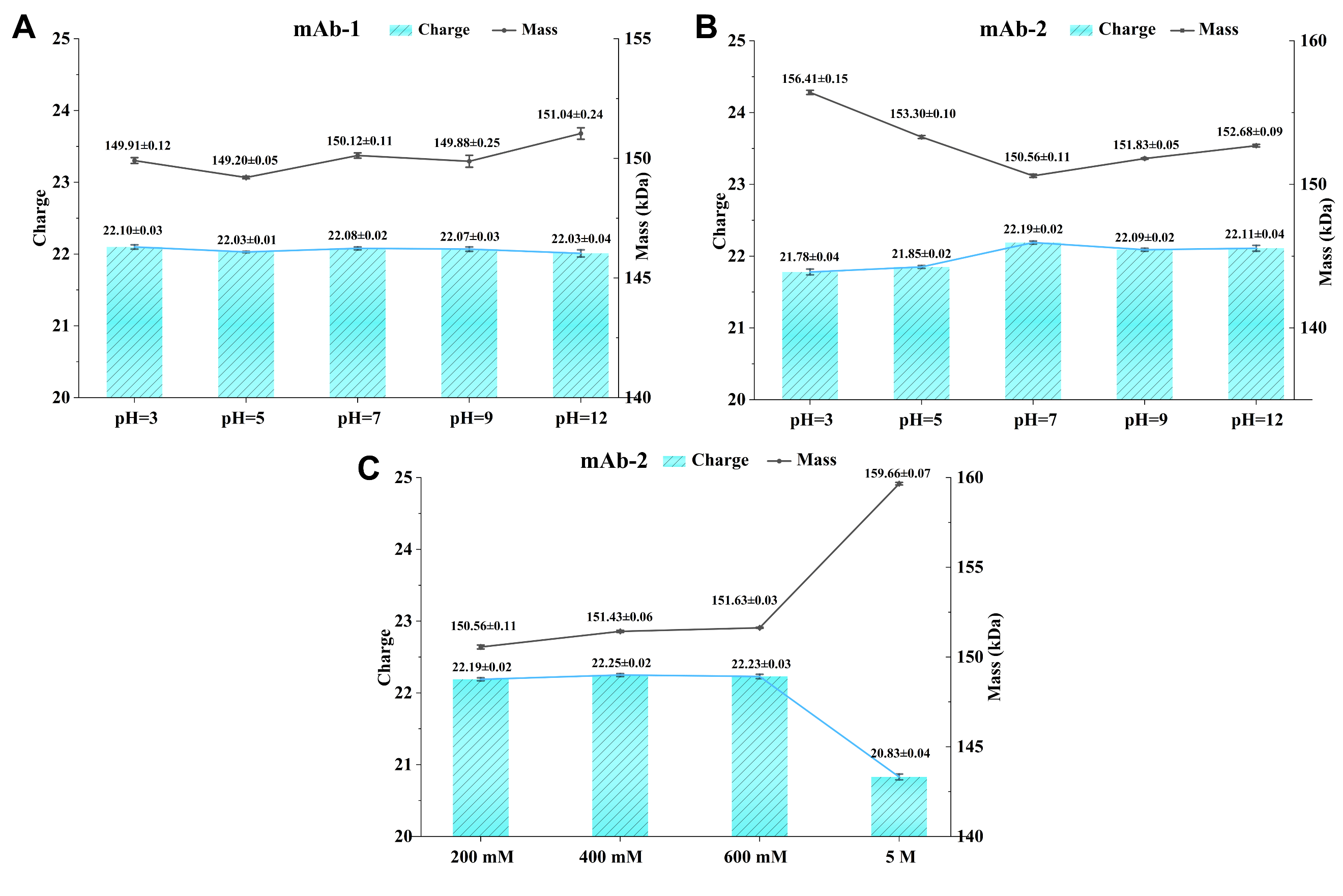

Figure S4. (A) mAb-1 in buffers at different pH values; (B) mAb-2 in buffers at different pH values; (C) mAb-2 in buffers at varying ammonium acetate concentrations. All datasets were derived from Gaussian fitting of raw mass and charge measurements. Bar charts represent charge state distributions, while scatter plots represent mass distributions. mAb-2 is known to be more sensitive to solution pH than mAb-1, and indeed showed larger fluctuation of measured mass and charge, suggesting the CDMS measurement is responsive to the mAb’s physicochemical properties.

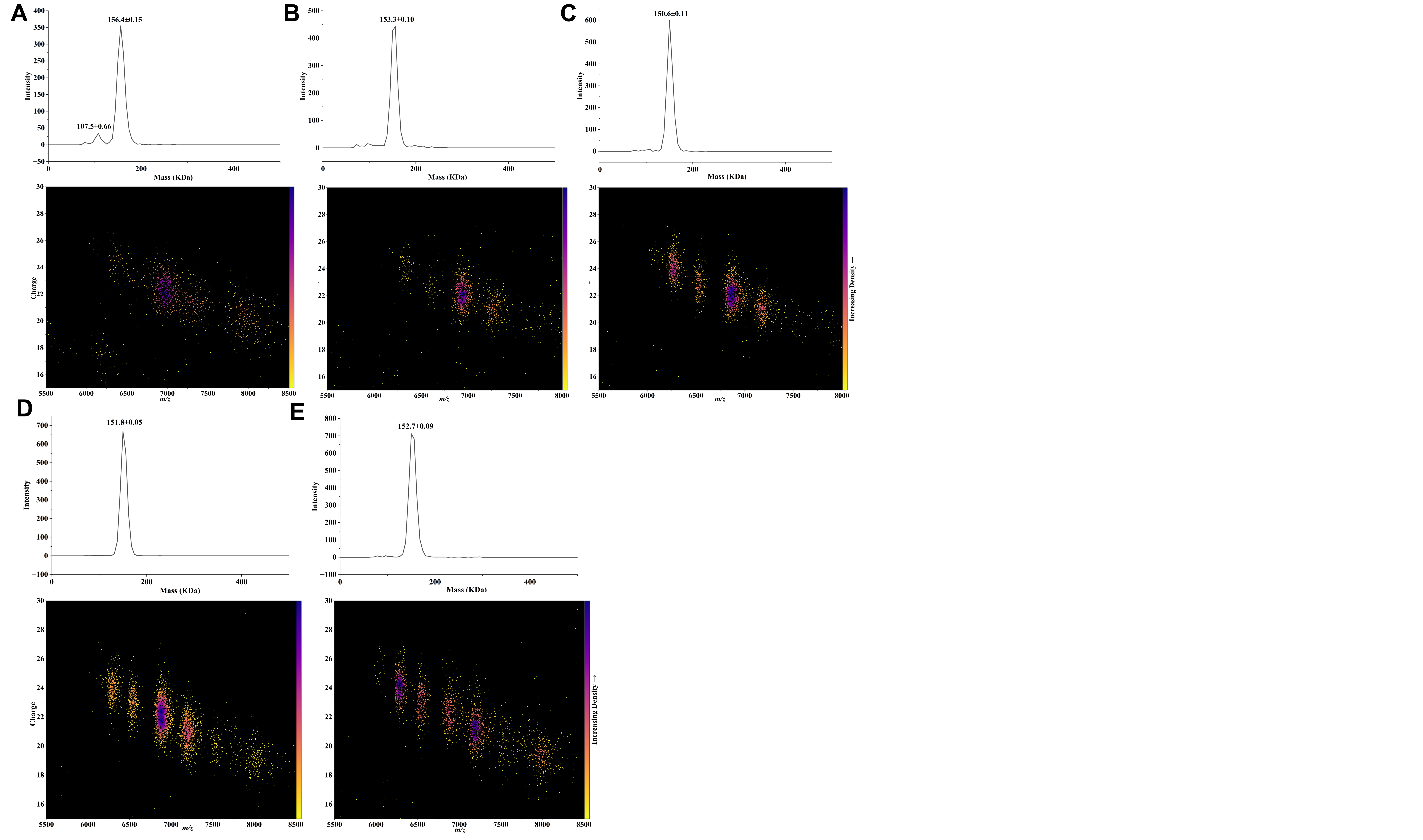

Figure S5. Native charge detection mass spectrometry of mAb-2 at different pH conditions. (A) pH=3, (B) pH=5, (C) pH=7.2, (D) pH=9, and (E) pH=12. For each pH, Top: Mass spectrum; Bottom: *m/z* vs. charge 2D point density plot. The mass values in the mass spectrum were obtained after Gaussian fitting.

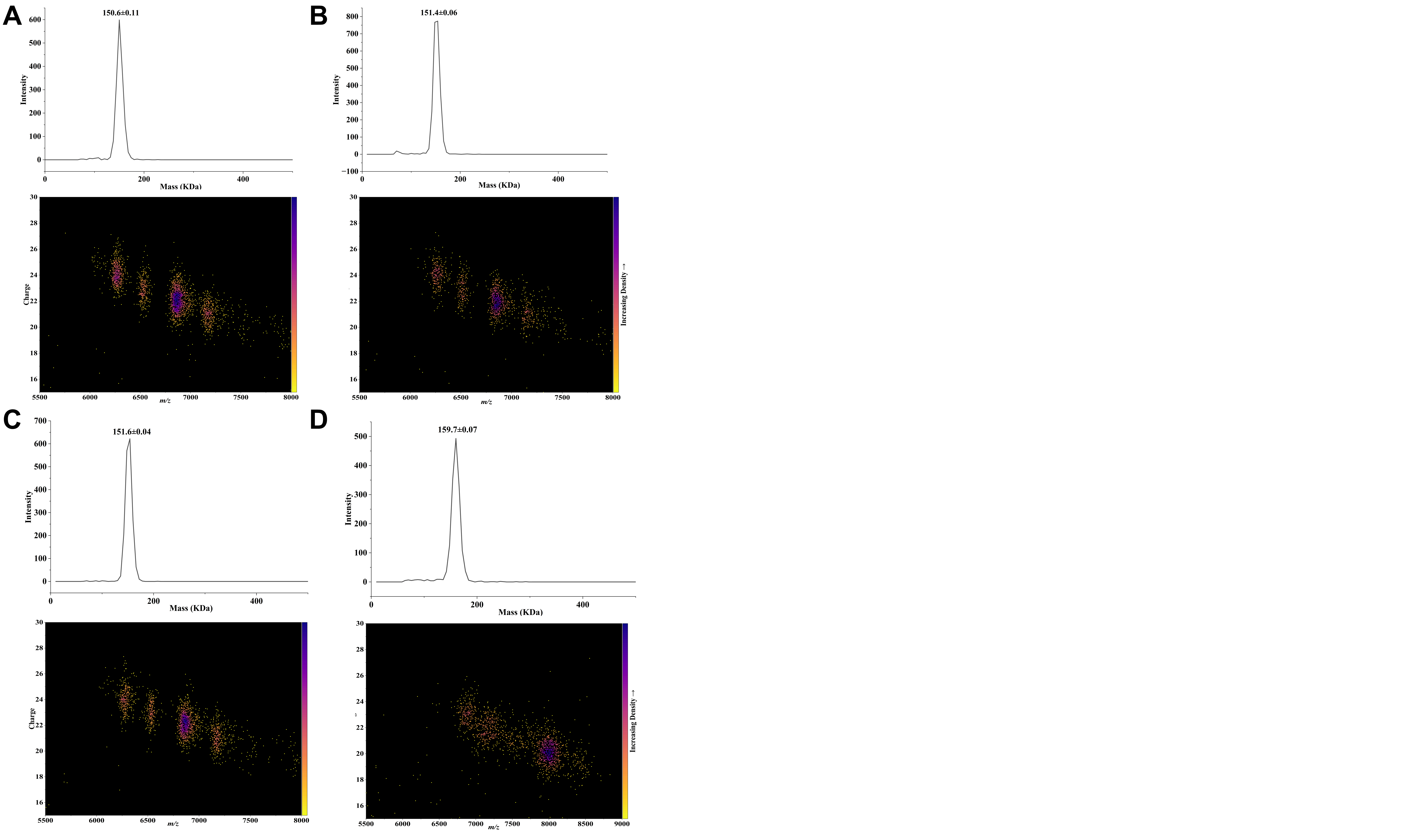

Figure S6. Native charge detection mass spectrometry of mAb-2 at different ammonium acetate concentration. (A) 200 mM, (B) 400 mM, (C) 600 mM, and (D) 5 M. For each concentration, Top: Mass spectrum; Bottom: *m/z* vs. charge 2D point density plot. The mass values in the mass spectrum were obtained after Gaussian fitting.

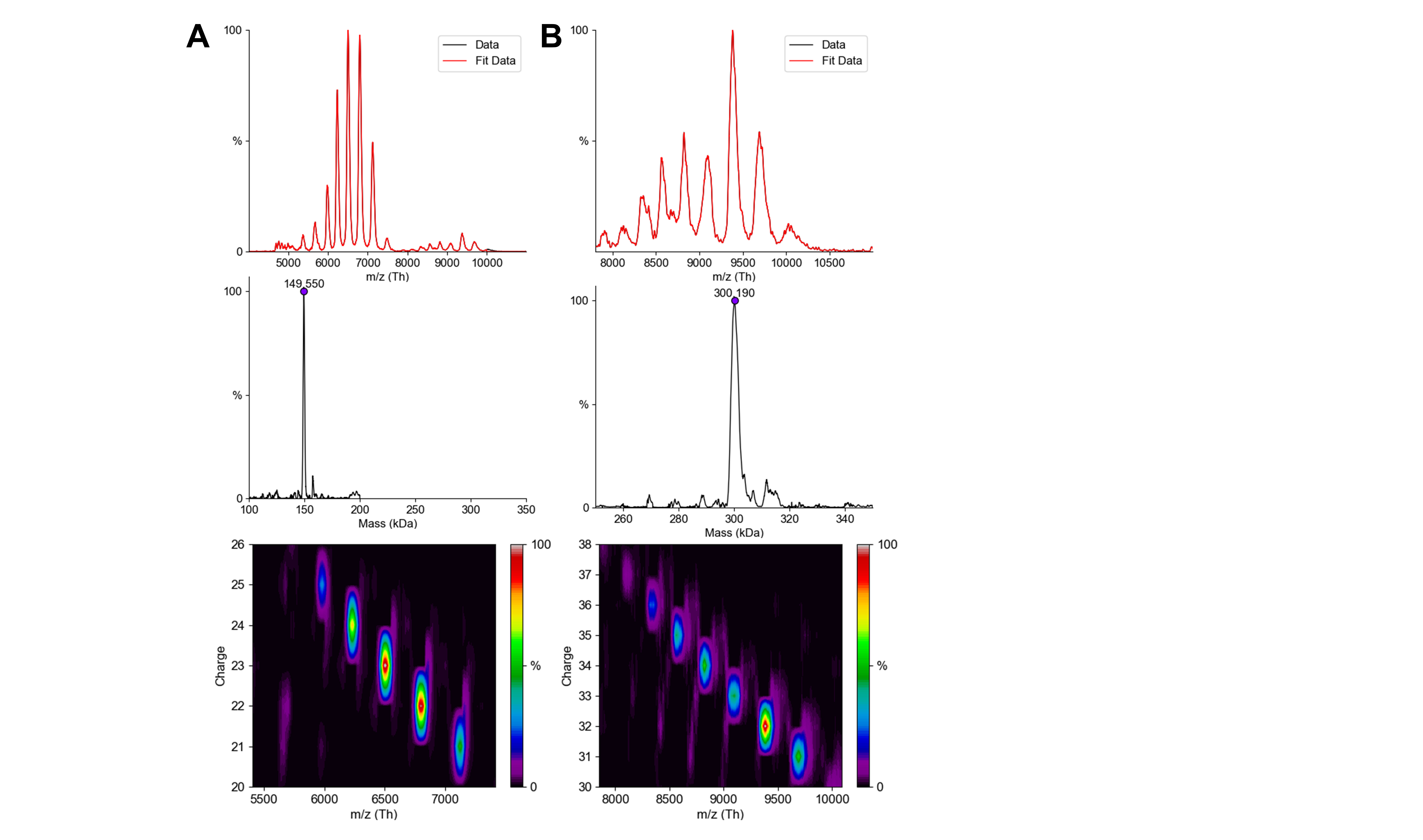

Figure S7. Native mass spectrometry of the full sample of mAb-1 after stress. (A) Monomer, and (B) Dimer. Top: *m/z* spectrum; Mid: Mass spectrum; Bottom: *m/z* vs. charge 2D plot. All data was obtained by unidec through convolution.

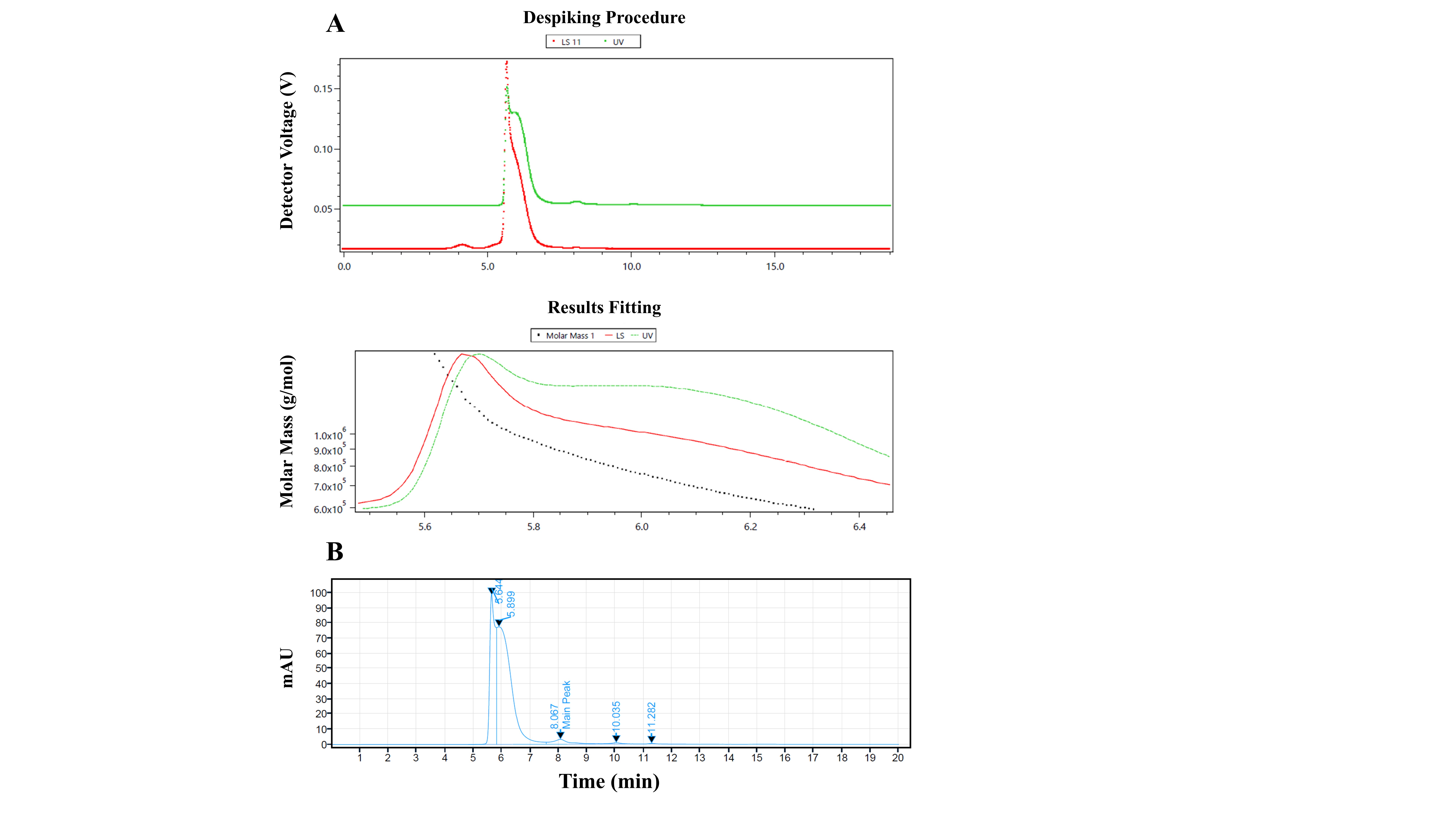

Figure S8. SEC and SEC-MALS chromatograms of the H1 fraction. (A) SEC-MALS chromatogram for molecular weight determination, the light scattering (LS) signal is shown in red and the ultraviolet (UV) absorbance signal in green. (B) SEC chromatogram used for purity quantification.

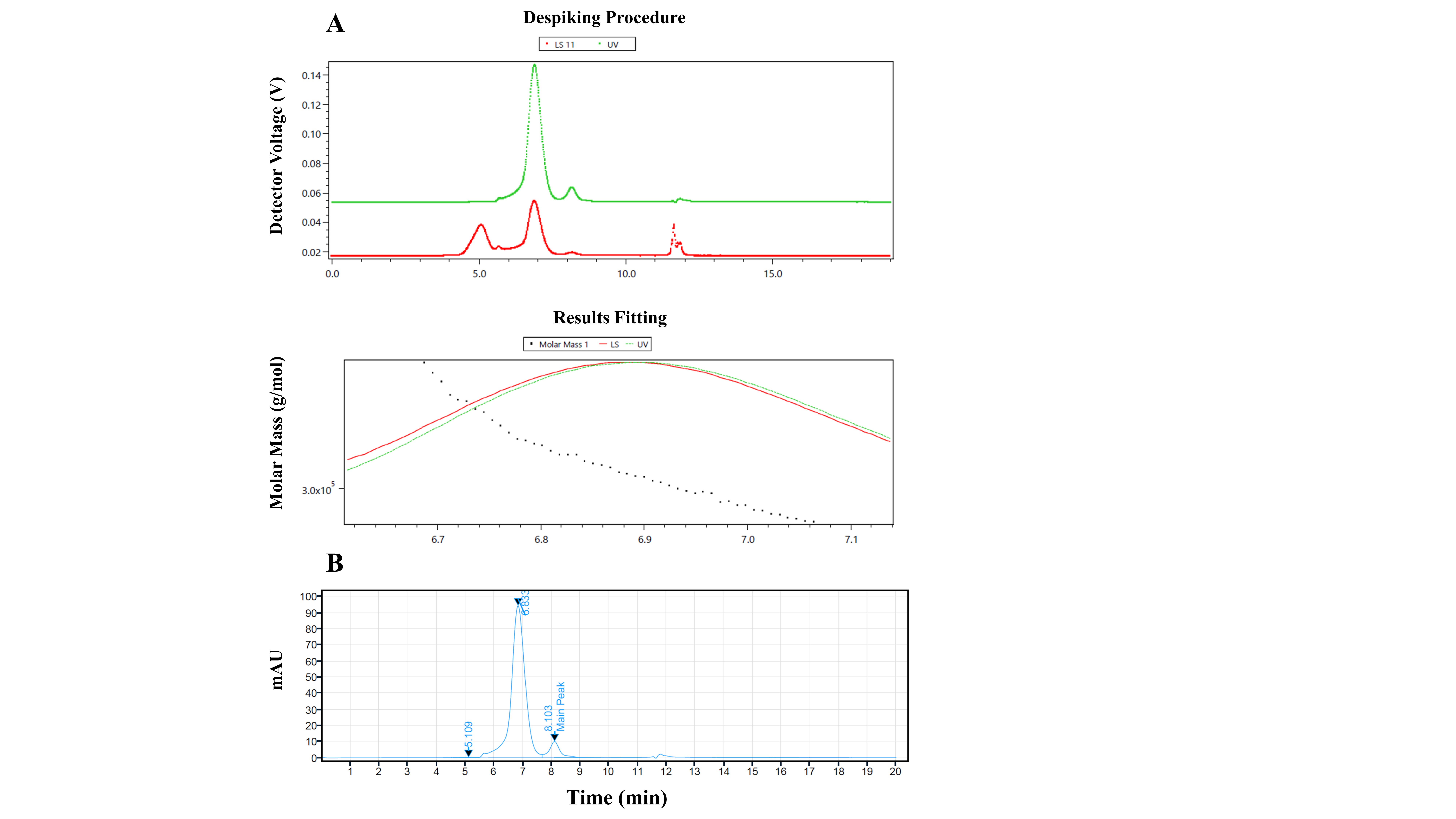

Figure S9. SEC and SEC-MALS chromatograms of the H2 fraction. (A) SEC-MALS chromatogram for molecular weight determination, LS signal is shown in red and the UV absorbance signal in green. (B) SEC chromatogram used for purity quantification.

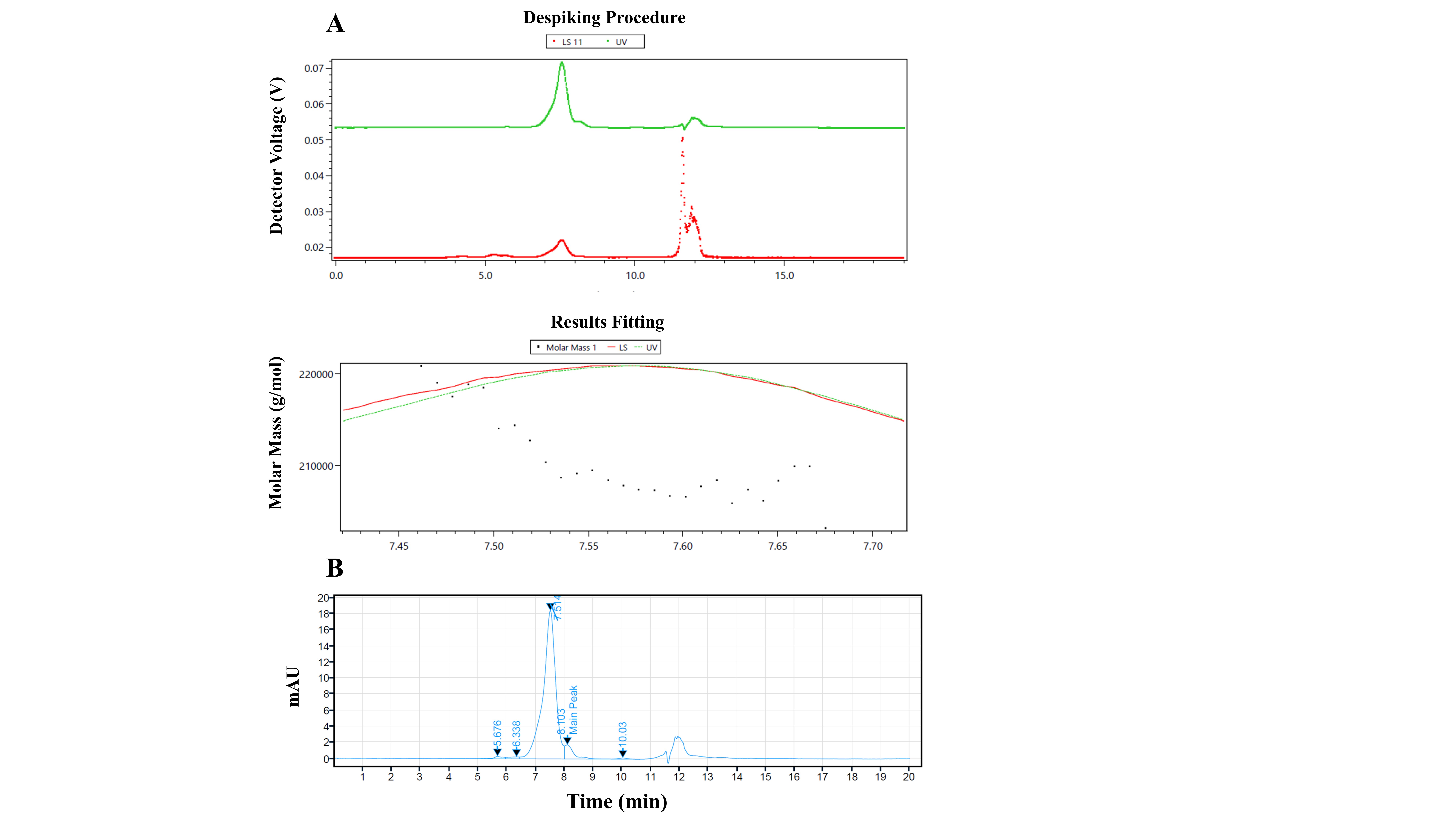

Figure S10. SEC and SEC-MALS chromatograms of the H3 fraction. (A) SEC-MALS chromatogram for molecular weight determination, LS signal is shown in red and the UV absorbance signal in green. (B) SEC chromatogram used for purity quantification.

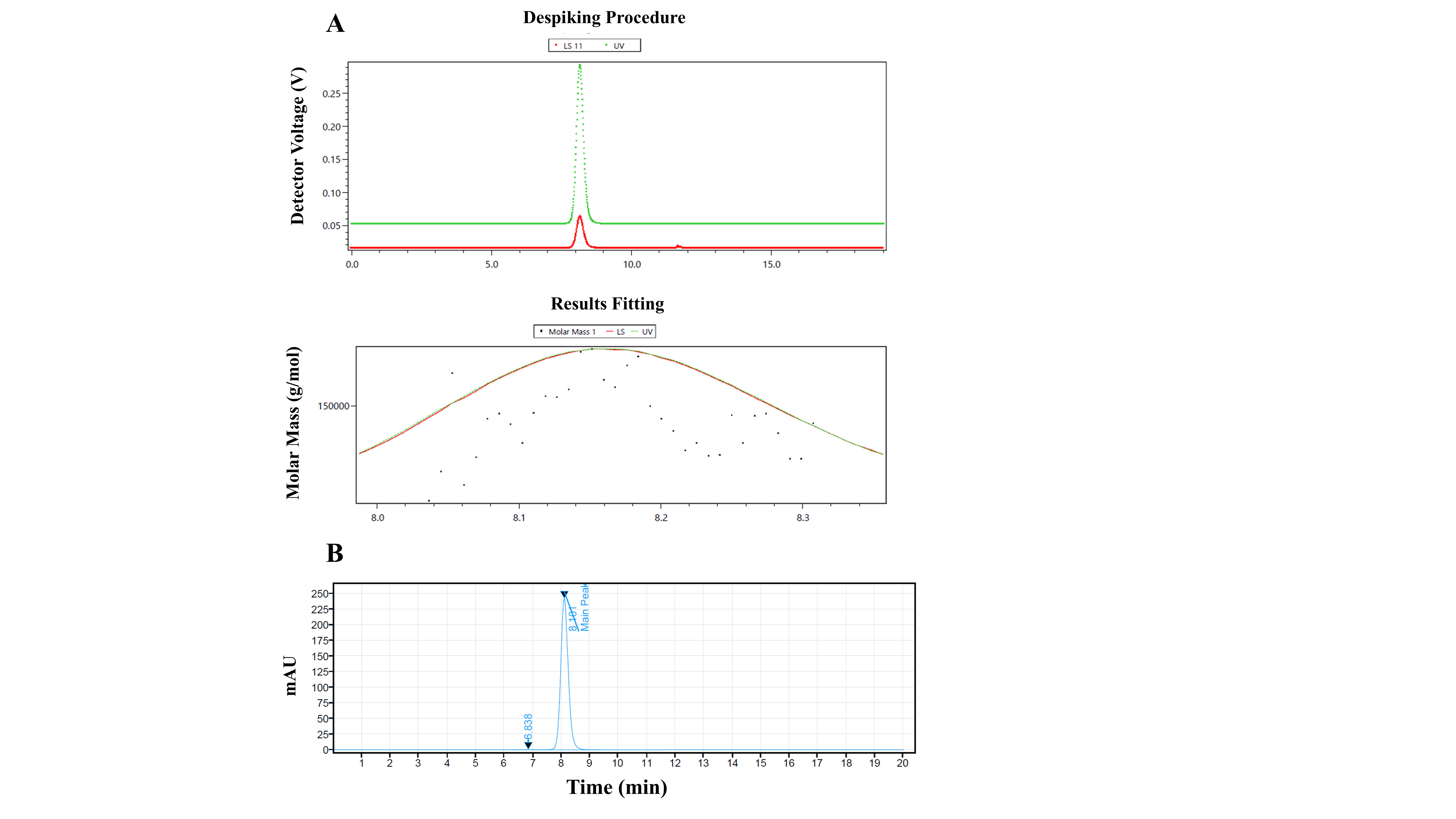

Figure S11. SEC and SEC-MALS chromatograms of the M fraction. (A) SEC-MALS chromatogram for molecular weight determination, LS signal is shown in red and the UV absorbance signal in green. (B) SEC chromatogram used for purity quantification.

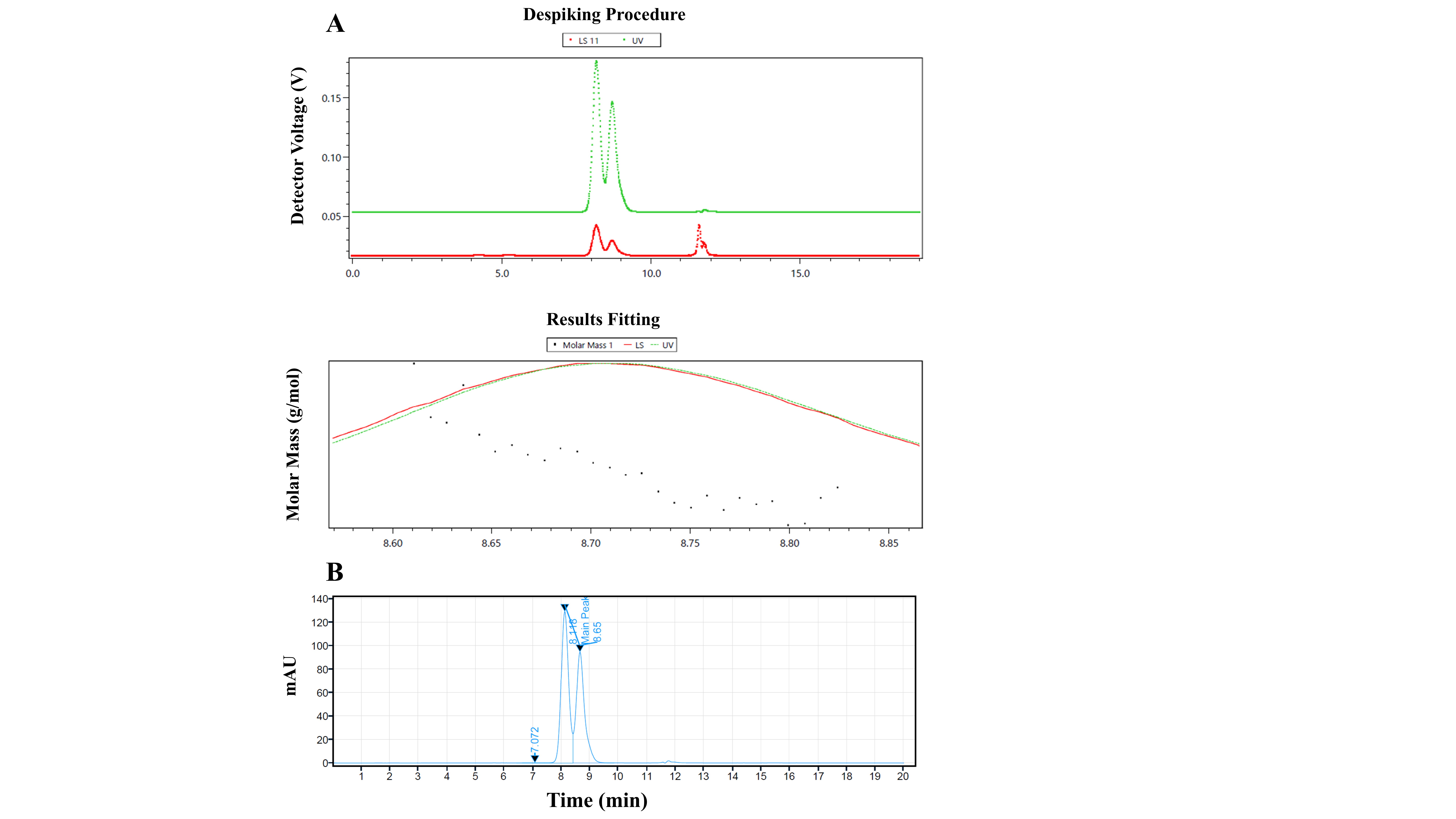

Figure S12. SEC and SEC-MALS chromatograms of the L1 fraction. (A) SEC-MALS chromatogram for molecular weight determination, LS signal is shown in red and the UV absorbance signal in green. (B) SEC chromatogram used for purity quantification.

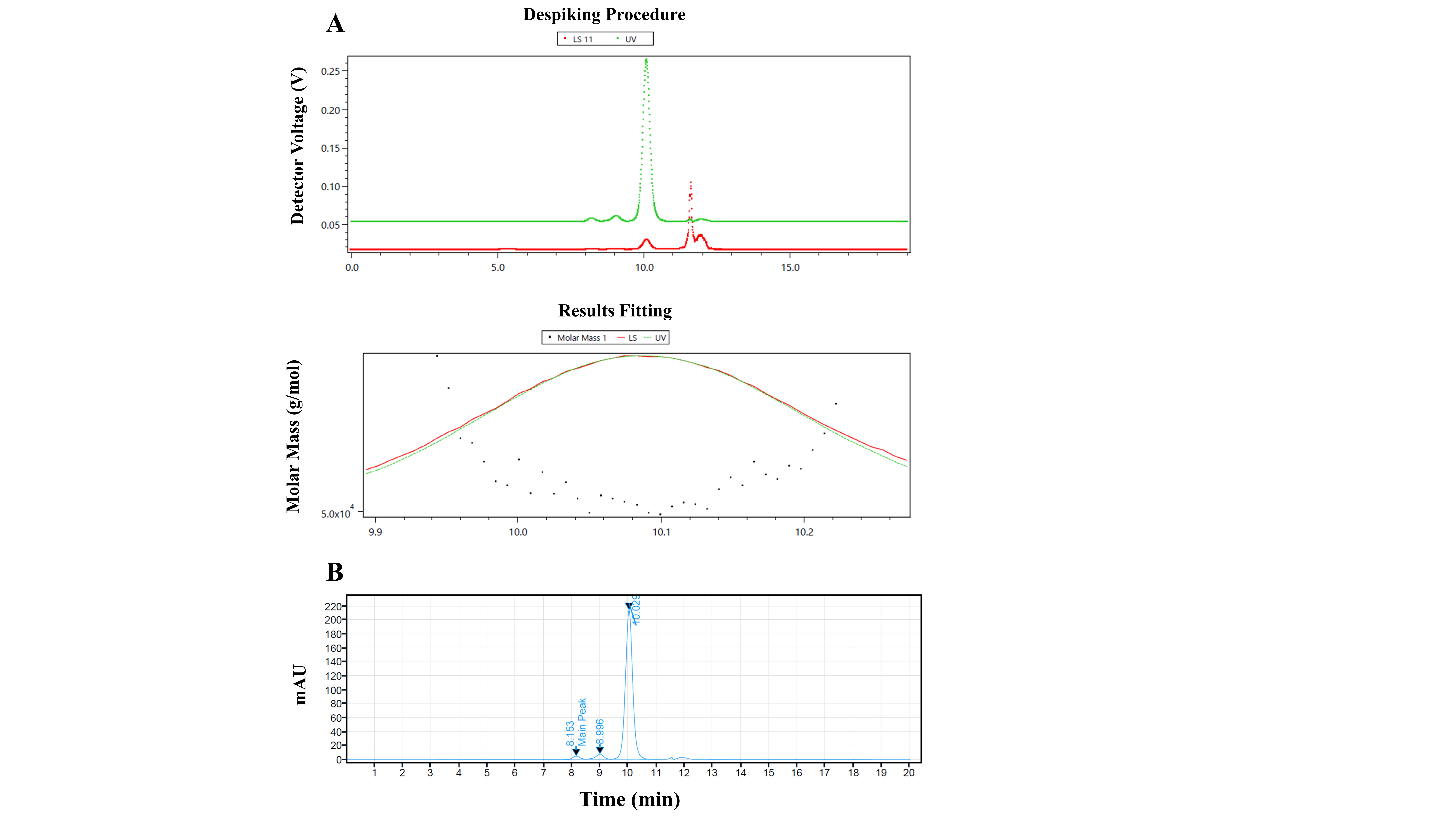

Figure S13. SEC and SEC-MALS chromatograms of the L2 fraction. (A) SEC-MALS chromatogram for molecular weight determination, LS signal is shown in red and the UV absorbance signal in green. (B) SEC chromatogram used for purity quantification.

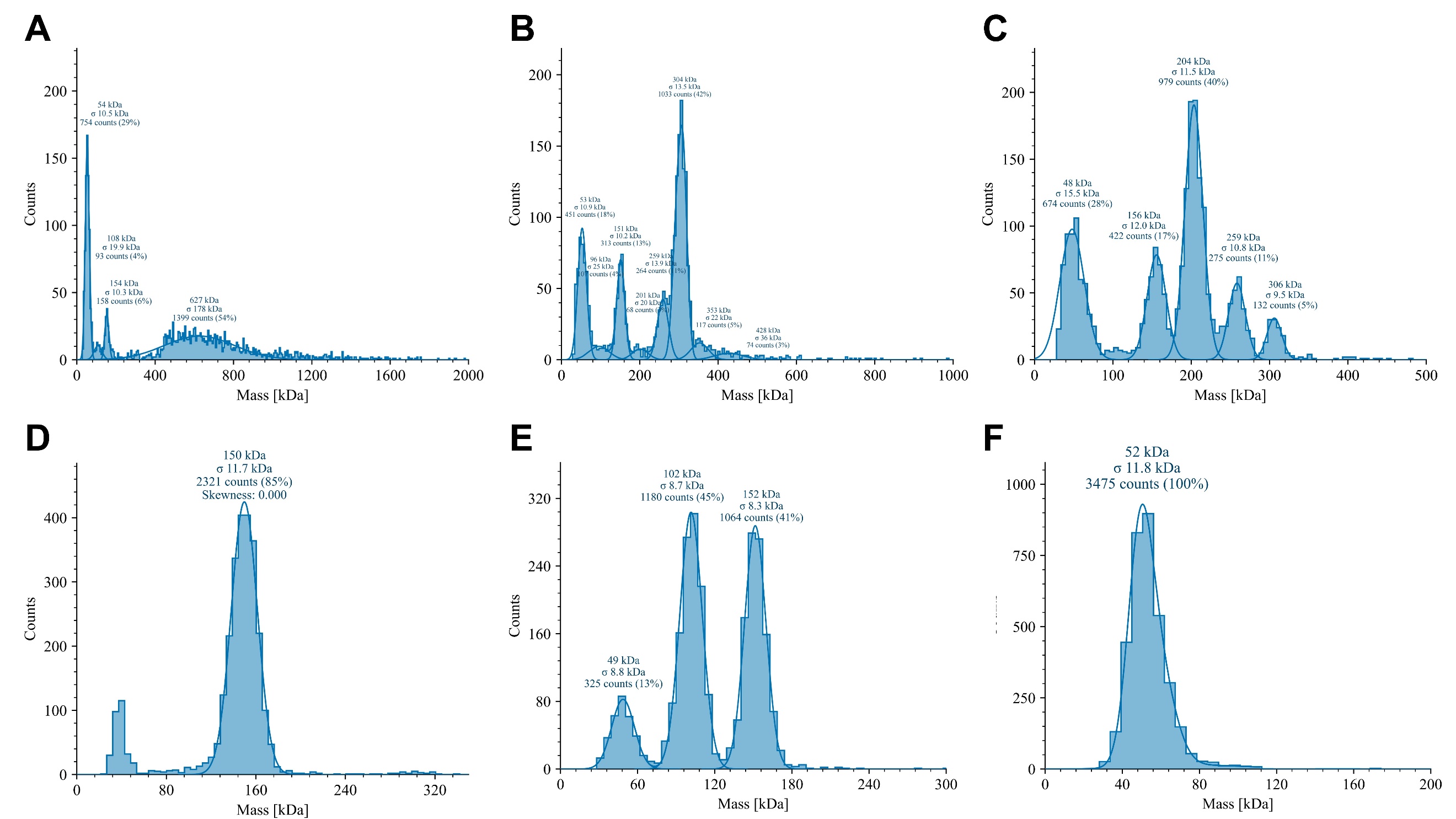

Figure S14. MP detection spectra for the six fractions. (A): H1, (B): H2, (C): H3, (D): M, (E): L1, and (F): L2. All data was processed using Refeyn DiscoverMP^TM^ software.

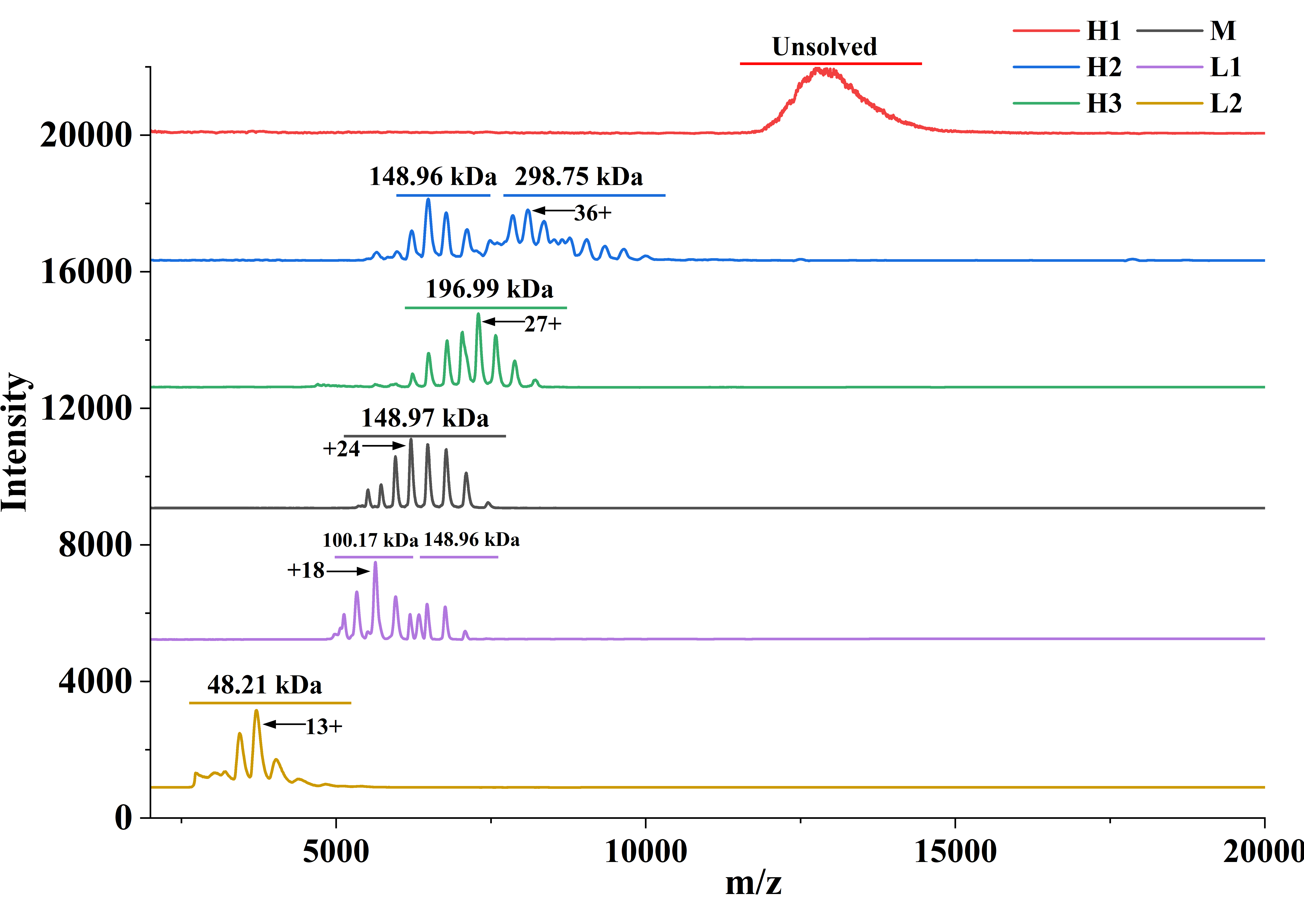

Figure S15. Smoothed native mass spectrometry spectra of the six SEC fractions. Deconvoluted mass values are annotated for fractions with resolvable ESI charge‑state distributions. Charge states of the H1 fraction could not be sufficiently resolved for deconvolution. The low *m/z* were filtered out in the method for qualitative analysis and improved sensitivity on high *m/z*. All mass and charge values were derived from raw data processed with UniDec software.

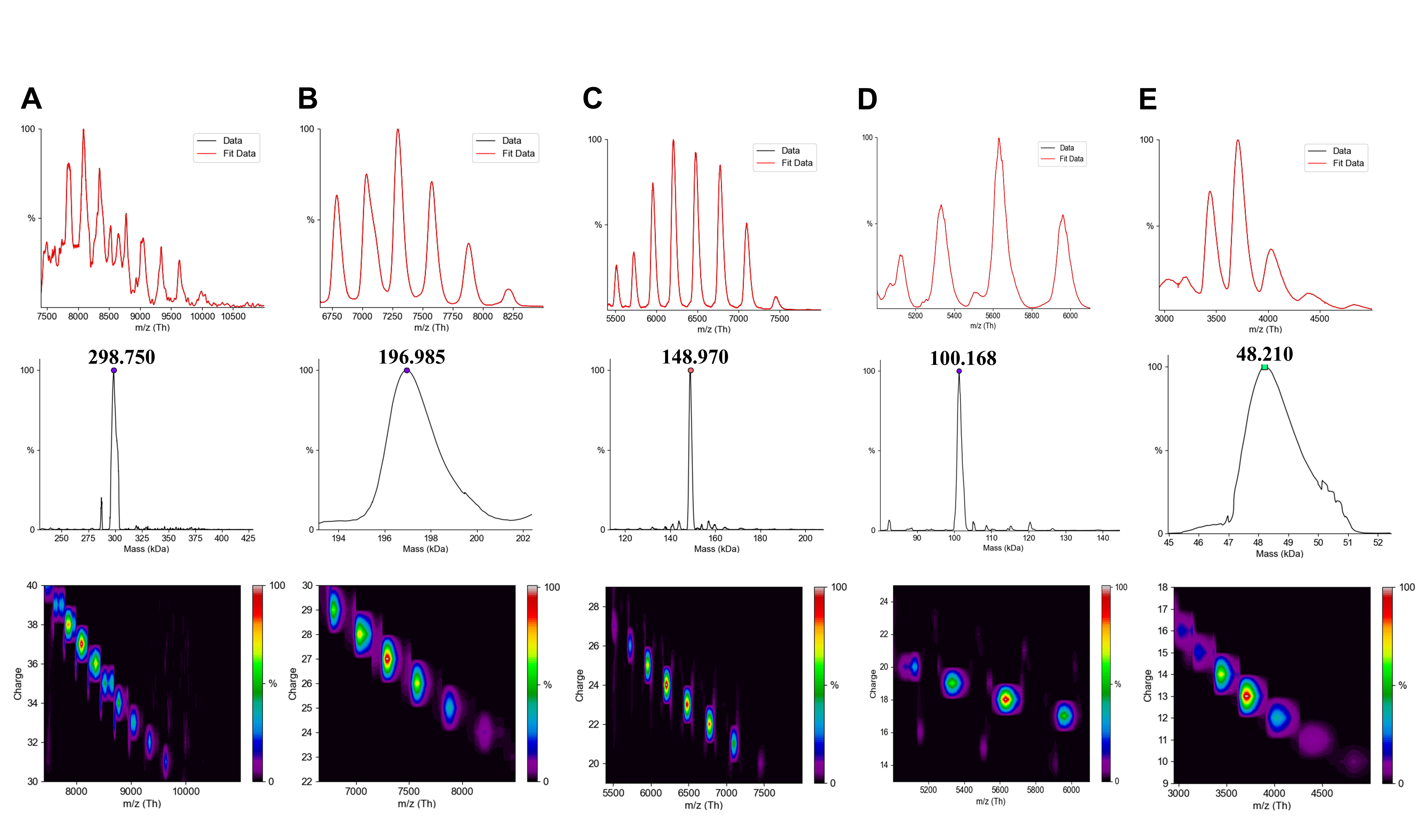

Figure S16. Deconvoluted native MS spectra of the indicated fractions. Deconvolution of H1 was not possible owing to its heterogeneity. Panels (A), (B), (C), (E), and (F) show H2, H3, M, L1, and L2, respectively. Only the main peaks of each fraction are shown. Top: *m/z* spectrum; Mid: Mass spectrum; Bottom: *m/z* vs. charge 2D plot. All mass and charge values were derived from raw data processed with UniDec software.

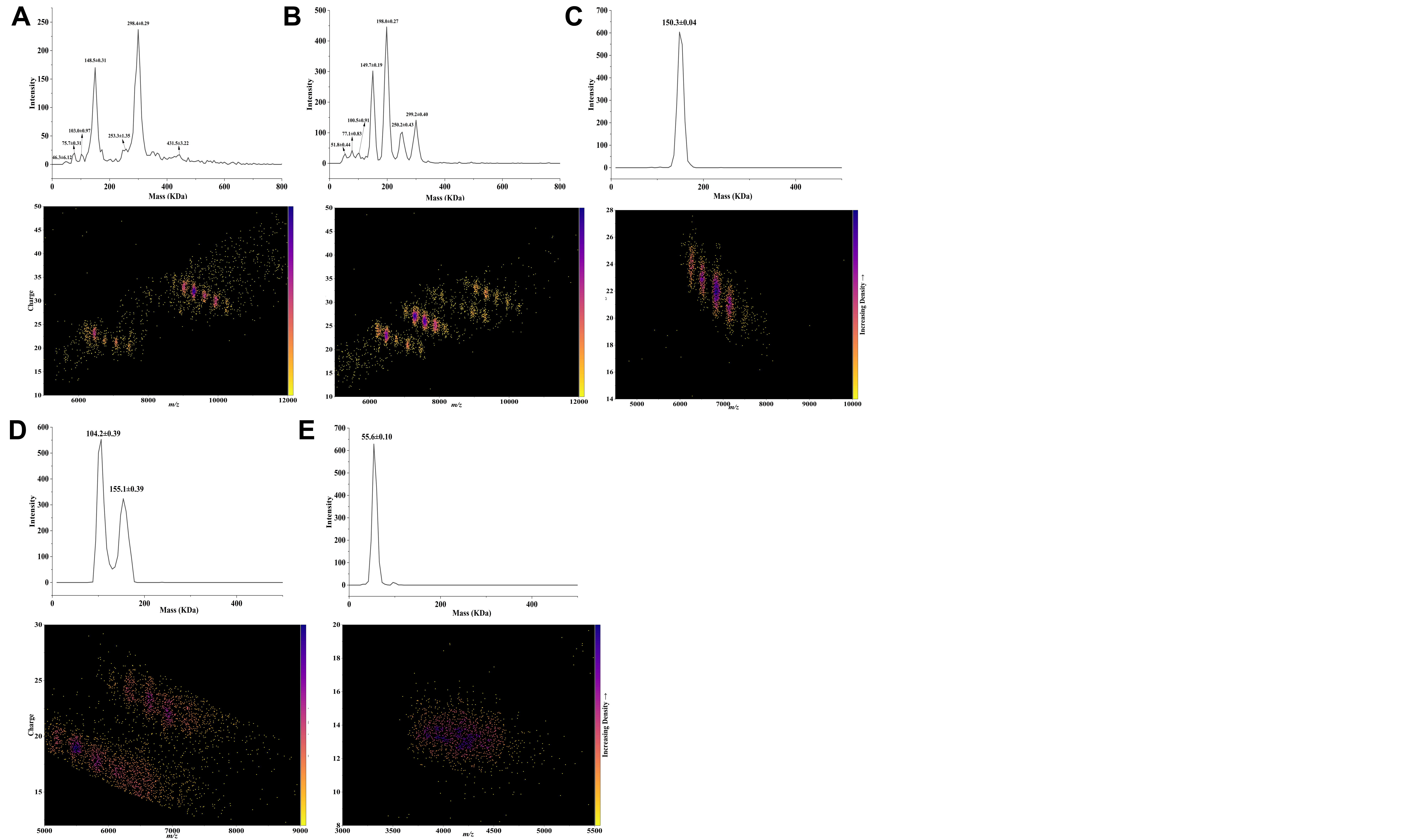

Figure S17. CDMS detection spectra for the six fractions without H1. (A): H2, (B): H3, (C): M, (D): L1, and (E): L2. For each fraction, Top: Mass spectrum; Bottom: *m/z* vs. charge 2D point density plot. The mass values in the mass spectrum were obtained after Gaussian fitting.

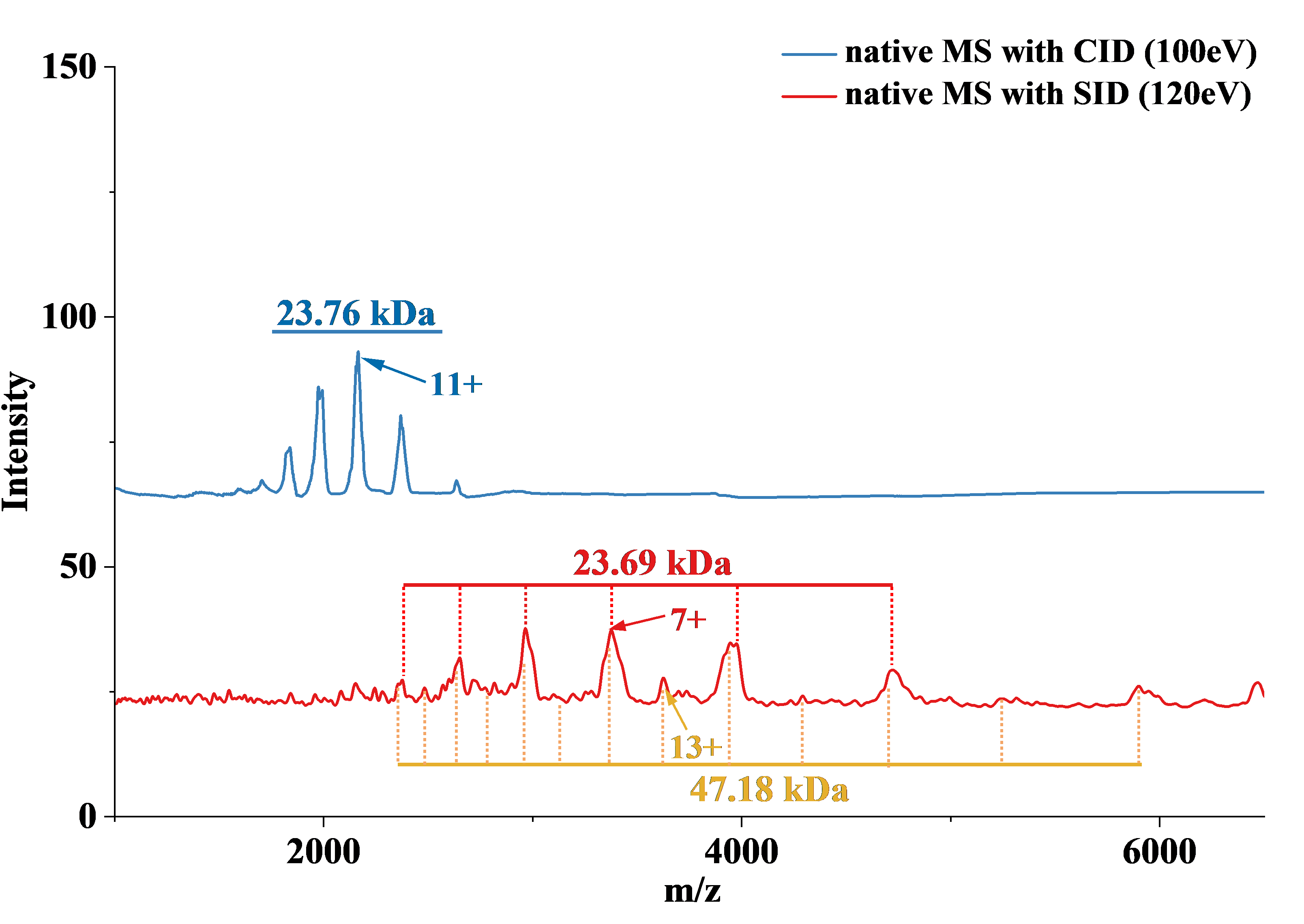

Figure S18. Native MS characterization of oligomer composition in the H3 fraction. The spectra were obtained using surface-induced dissociation (SID, 120 eV, red trace) and collision-induced dissociation (CID, 100 eV, blue trace), respectively.

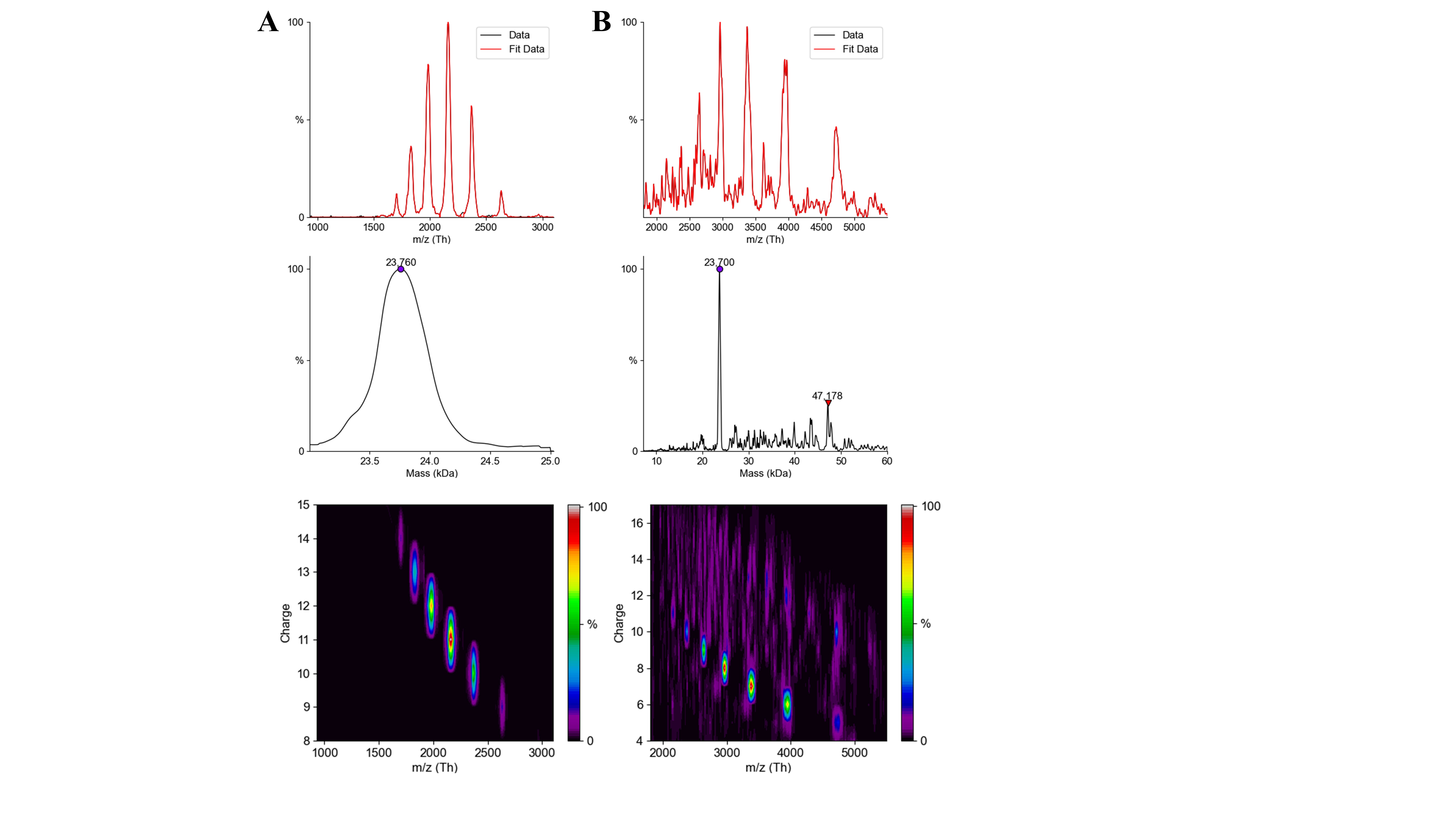

Figure S19. Native mass spectrometry of the H3 fraction. (A) CID, and (B) SID. Top: *m/z* spectrum; Mid: Mass spectrum; Bottom: *m/z* vs. charge 2D plot. All mass and charge values were derived from raw data processed with UniDec software.

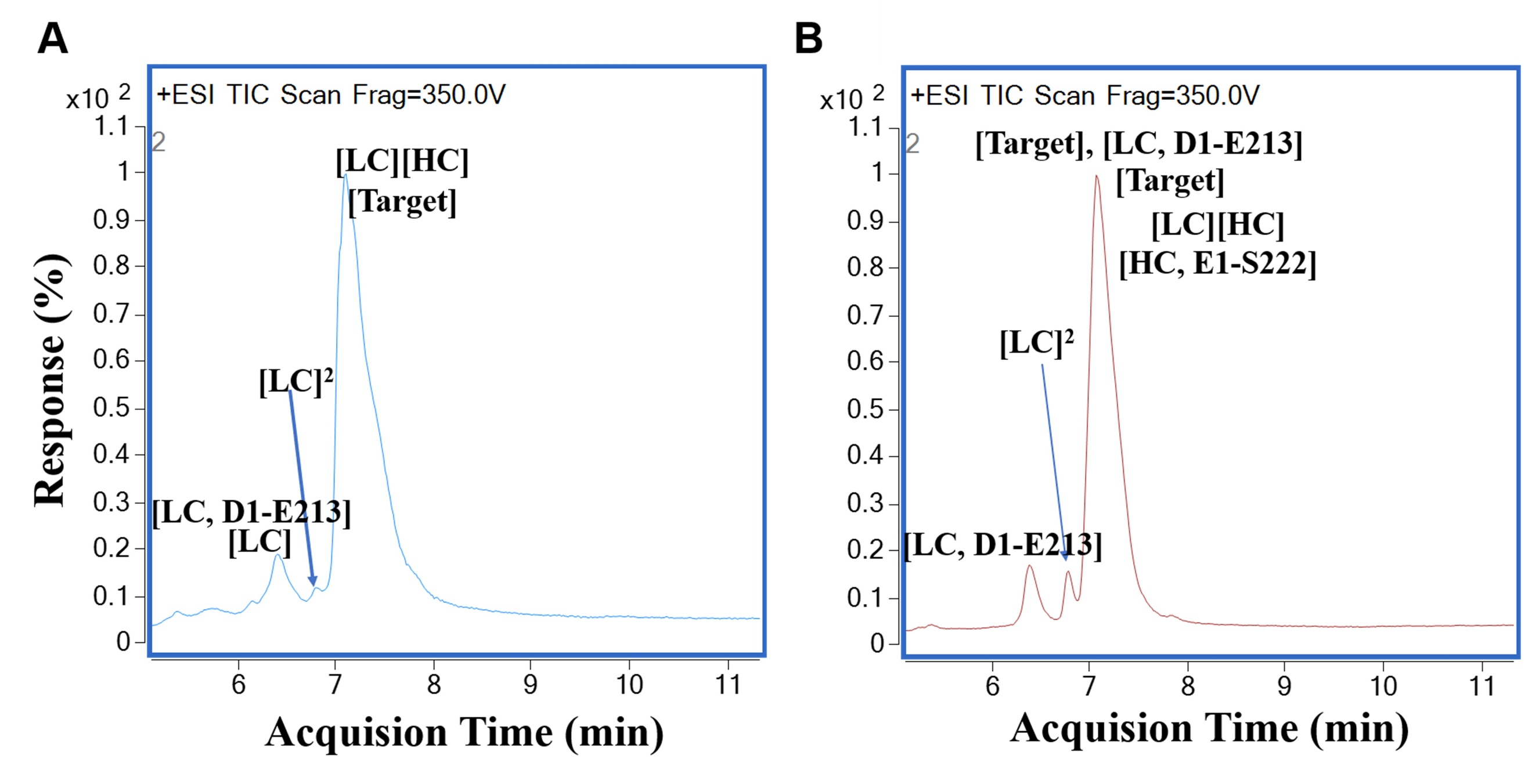

Figure S20. Characterization of oligomer composition in the H1 and H3 fractions. (A) RP LC-MS analysis of H1 fraction. Detailed data is shown in Table S6; (B) RP LC-MS analysis of H3 fraction. The total ion chromatogram (TIC) shows the separation of eluted species, with identified components and chains labeled. Detailed data is shown in Table S7.

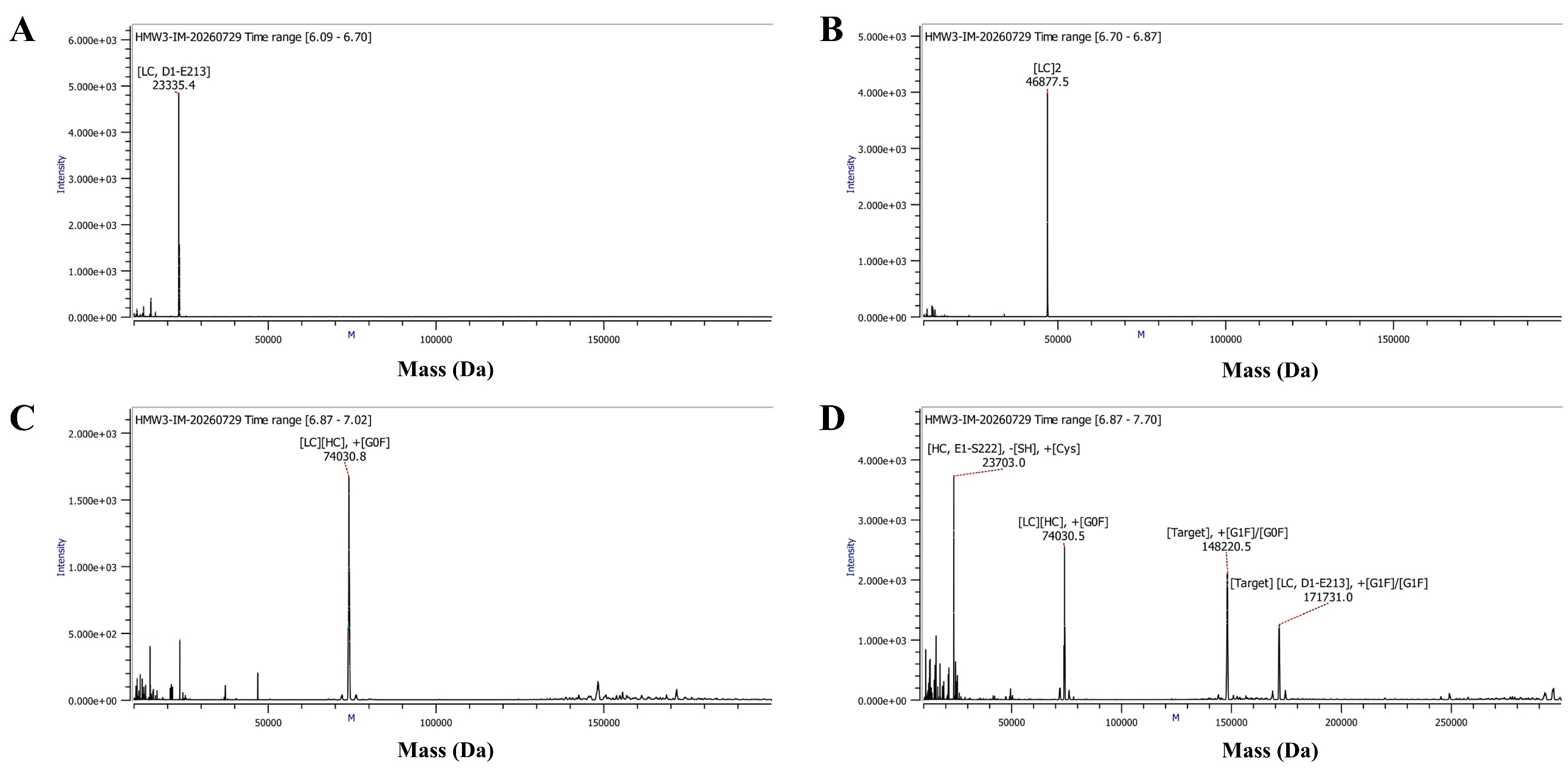

Figure S21. Deconvoluted mass spectra of the H3 fraction from RP LC-MS analysis at different retention time windows: (A) In the 6.09–6.70 min window; (B) In the 6.70–6.87 min window; (C) In the 6.87–7.02 min window; (D) In the 6.87–7.70 min window.

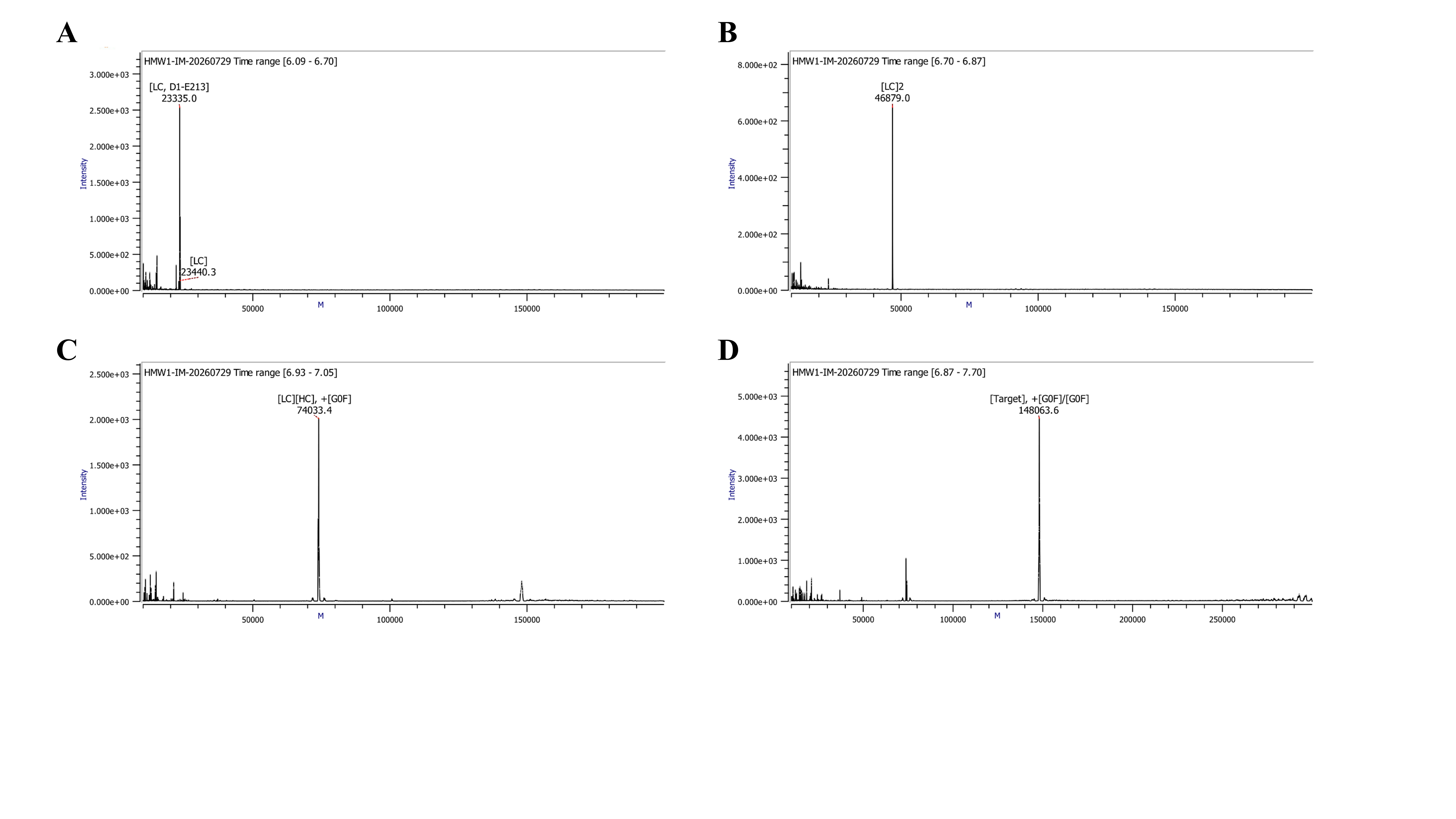

Figure S22. Deconvoluted mass spectra of the H1 fraction from RP LC-MS analysis at different retention time windows: (A) In the 6.09–6.70 min window; (B) In the 6.70–6.87 min window; (C) In the 6.87–7.02 min window; (D) In the 6.87–7.70 min window.

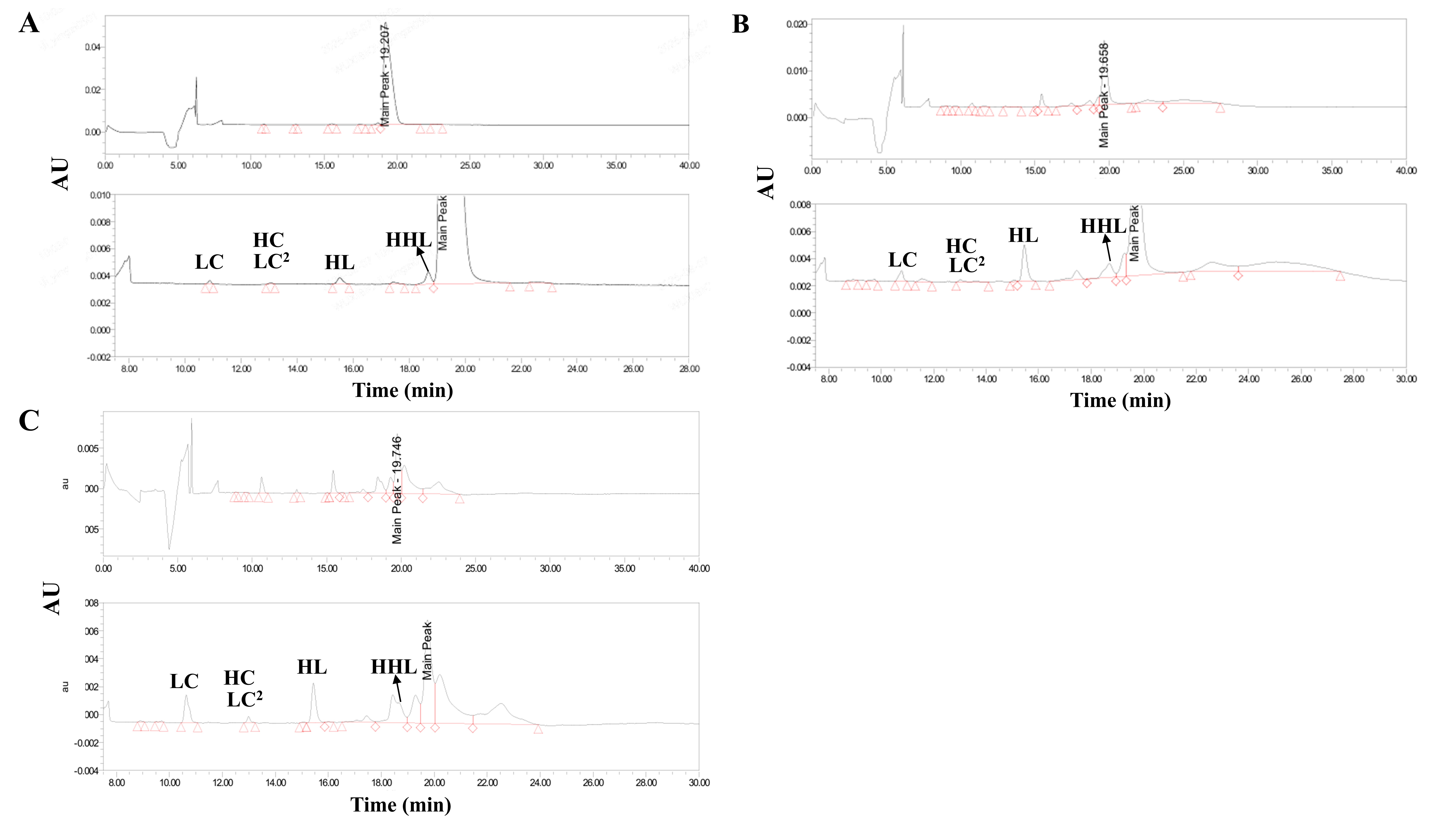

Figure S23. Non-reduced capillary electrophoresis-sodium dodecyl sulfate (nrCE-SDS) analysis of H1 and H3 fractions. (A) Intact mAb-1 control sample; (B) H1 fraction; (C) H3 fraction. For each panel, the upper trace presents the full view of the electropherogram, while the lower trace shows the zoomed-in region to highlight low-abundance species and minor peaks. The retention time positions of relevant peaks are marked with red open triangles. Species related to assembly, including light chain (LC), heavy chain (HC), light chain dimer (LC^2^), half-molecule (HL), and heavy-heavy-light chain (HHL), are labeled accordingly.

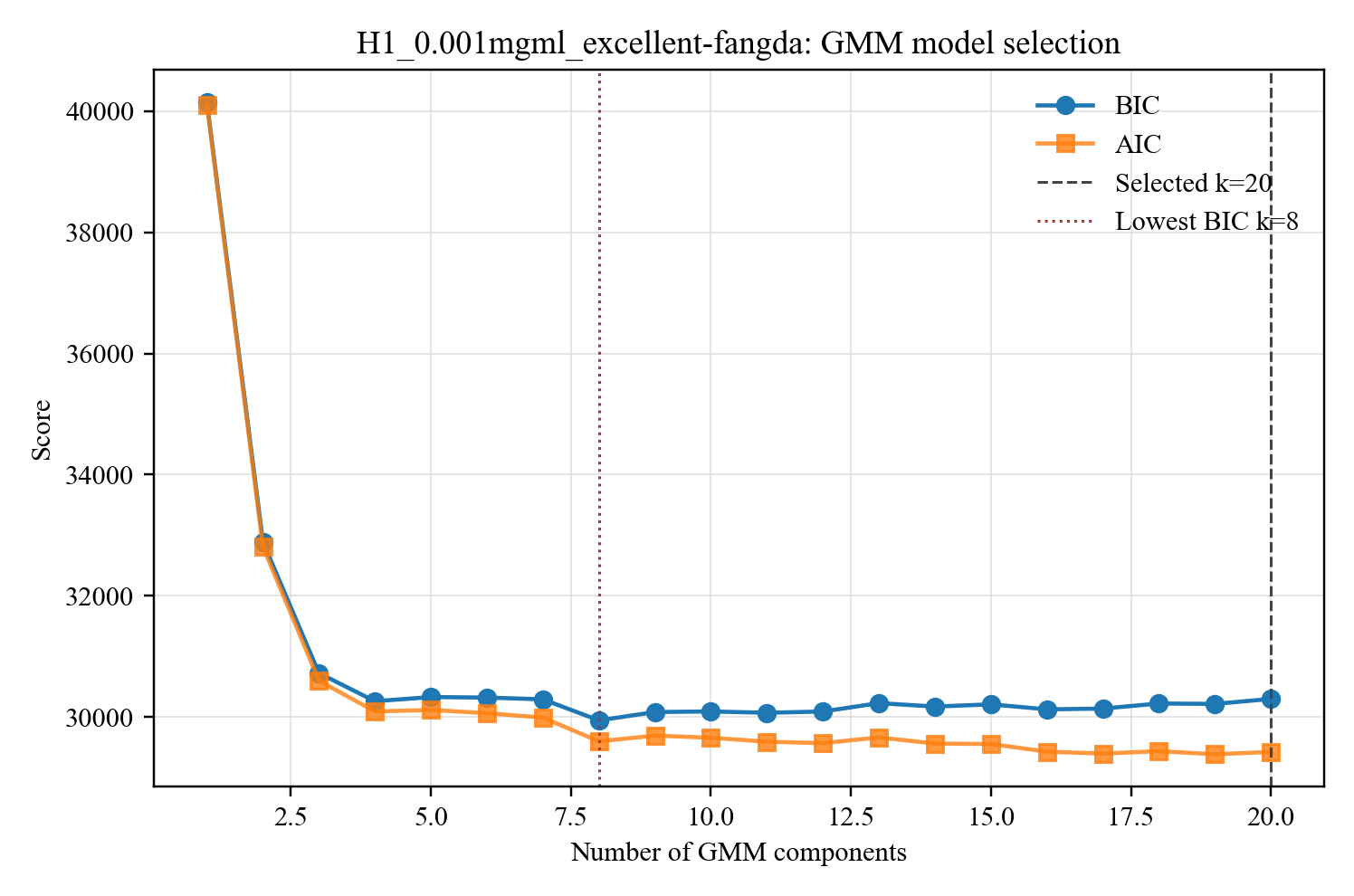

Figure S24. Selection of the number of components in the GMM based on AIC and BIC. The red dashed line indicates the number of GMM components (k = 8) corresponding to the minimum BIC value, while the black dashed line represents the number of components (k = 20) ultimately selected in this study.

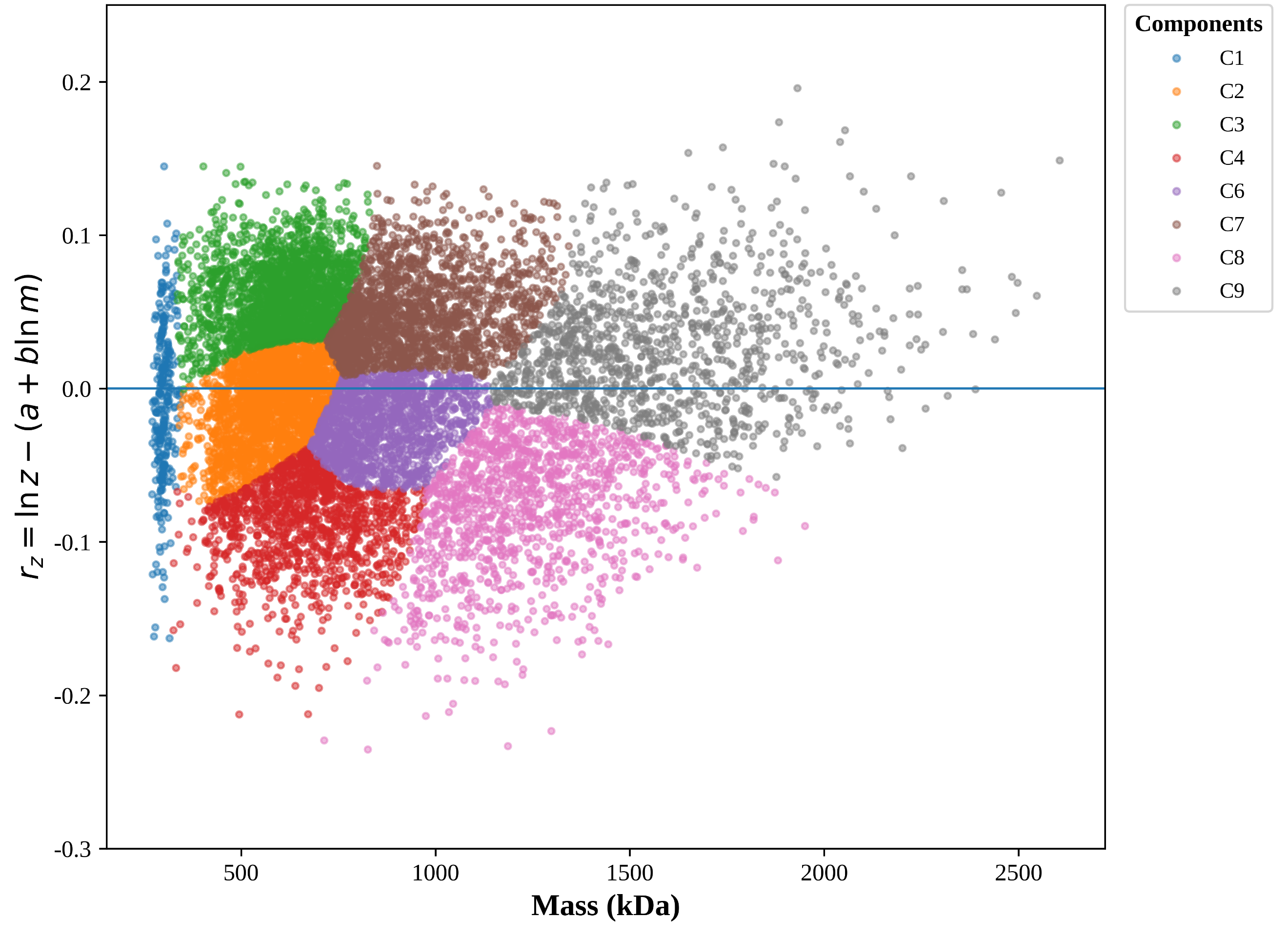

Figure S25. Mass-conditioned charge analysis of the major ion families in the H1 fraction. The expected mass-dependent charging relationship was first modeled as $\ln z=a+b\ln m$, where $m$and $z$denote the single-ion mass and charge, respectively. The mass-conditioned charge residual was subsequently calculated as $r_{z}=\ln z-\left( a+b\ln m \right)$, such that positive and negative residuals represent charging above and below that expected for ions of comparable mass, respectively. Gaussian mixture modeling was performed in the two-dimensional $\left( \ln m , r_{z} \right)$space. BIC analysis favored a nine-component model. For clarity, the minor C5 component, which accounted for approximately 0.74% of the fitted population and contained the high-residual tail, is omitted from the present visualization; the assignments of the remaining components were retained from the original nine-component model. Each point represents an individual ion and is colored according to its maximum-posterior GMM assignment. The substantial overlap among neighboring components indicates that these groups represent statistical ion families rather than unequivocally discrete molecular species.

### **Supplementary Tables:**

Table S1. The original formulation of two mAbs

| Serial numbers | Concentration (mg/mL) | Formulations |
| --- | --- | --- |
| mAb-1 | 50 | 20 mM His, 8% sucrose (pH 5.8) |
| mAb-2 | 20 | 100 mg/mL sucrose, 0.1 mg/mL polysorbate 80, 20 mM phosphate (pH 7.2) |

Table S2. Mass repeatability data from six consecutive measurements (n=6).

| Samples | Mass of monomer (kDa) | Mean (kDa) | RSD (%) | Mass of dimer (kDa) | Mean (kDa) | RSD (%) |
| --- | --- | --- | --- | --- | --- | --- |
| 1 | 68.36 | 68.49±0.15 | 0.21 | 136.31 | 136.30±0.64 | 0.47 |
| 2 | 68.29 |  |  | 135.33 |  |  |
| 3 | 68.46 |  |  | 135.76 |  |  |
| 4 | 68.63 |  |  | 136.72 |  |  |
| 5 | 68.45 |  |  | 136.65 |  |  |
| 6 | 68.72 |  |  | 137.04 |  |  |

Table S3. Mass inter-precision data of different data sets measured by different professionals (n=6).

| Samples | Mass of monomer (kDa) | Mean (kDa) | RSD (%) | Mass of dimer (kDa) | Mean (kDa) | RSD (%) |
| --- | --- | --- | --- | --- | --- | --- |
| 1 | 68.53 | 68.47±0.11 | 0.16 | 136.12 | 136.50±0.83 | 0.61 |
| 2 | 68.35 |  |  | 135.51 |  |  |
| 3 | 68.33 |  |  | 135.24 |  |  |
| 4 | 68.56 |  |  | 136.19 |  |  |
| 5 | 68.49 |  |  | 136.08 |  |  |
| 6 | 68.58 |  |  | 137.87 |  |  |

Table S4 Charge repeatability data from six consecutive measurements (n=6).

| Samples | Charge of monomer | Mean | RSD (%) | Charge of dimer | Mean | RSD (%) |
| --- | --- | --- | --- | --- | --- | --- |
| 1 | 14.79 | 14.75±0.02 | 0.16 | 21.18 | 21.21±0.04 | 0.21 |
| 2 | 14.74 |  |  | 21.21 |  |  |
| 3 | 14.72 |  |  | 21.18 |  |  |
| 4 | 14.76 |  |  | 21.28 |  |  |
| 5 | 14.77 |  |  | 21.24 |  |  |
| 6 | 14.76 |  |  | 21.17 |  |  |

Table S5. Charge inter-precision data of different data sets measured by different professionals (n=6).

| Samples | Charge of monomer | Mean | RSD (%) | Charge of dimer | Mean | RSD (%) |
| --- | --- | --- | --- | --- | --- | --- |
| 1 | 14.76 | 14.76±0.03 | 0.20 | 21.17 | 21.20±0.08 | 0.37 |
| 2 | 14.75 |  |  | 21.12 |  |  |
| 3 | 14.75 |  |  | 21.16 |  |  |
| 4 | 14.72 |  |  | 21.13 |  |  |
| 5 | 14.81 |  |  | 21.31 |  |  |
| 6 | 14.78 |  |  | 21.28 |  |  |

Table S6. Analysis of the compositional data for the H3 fraction by RP LC-MS. [SH] refers to the conversion of cysteine to dehydroalanine. [Cys] refers to cysteine. G0F and G1F refer to N-glycans. Target represents the monomer of mAb-1.

| Species | Modification | Theoretical Mass (Da) | Measured Mass (Da) | Mass Error (ppm) | Mass Difference (Da) |
| --- | --- | --- | --- | --- | --- |
| [LC, D1-E213] | / | 23,335.8 | 23,335.4 | -14.5 | -0.3 |
| [HC, E1-S222] | -[SH], +[Cys] | 23,703.4 | 23,703.0 | -14.9 | -0.4 |
| [LC]^2^ | / | 46,875.8 | 46,877.5 | 36.5 | 1.7 |
| [LC][HC] | +[G0F] | 74,028.3 | 74,030.5 | 29.8 | 2.2 |
| [Target] | +[G1F]/[G0F] | 148,218.7 | 148,220.5 | 12.3 | 1.8 |
| [Target][LC, D1-E213] | +[G1F]/[G1F] | 171,716.6 | 171,731.0 | 83.8 | 14.4 |

Table S7. Analysis of the compositional data for the H1 fraction by RP LC-MS. [SH] refers to the conversion of cysteine to dehydroalanine. [Cys] refers to cysteine. G0F and G1F refer to N-glycans. Target represents the monomer of mAb-1. Mass Error = (Measured mass - Theoretical mass)/ Theoretical mass × 10⁶. The results were calculated with the raw data and reported with 1 decimal place.

| Species | Modification | Theoretical Mass (Da) | Measured Mass (Da) | Mass Error (ppm) | Mass Difference (Da) |
| --- | --- | --- | --- | --- | --- |
| [LC, D1-E213] | / | 23,335.8 | 23,335.0 | -31.3 | -0.7 |
| [LC] | / | 23,438.9 | 23,440.3 | 61.4 | 1.4 |
| [LC]^2^ | / | 46,875.8 | 46,879.0 | 69.0 | 3.2 |
| [LC][HC] | +[G0F] | 74,028.3 | 74,033.4 | 68.4 | 5.1 |
| [Target] | +[G0F]/[G0F] | 148,056.6 | 148,063.6 | 47.1 | 7.0 |

Table S8. Summary of GMM-resolved subpopulations in the H1 fraction. Tentative assignments were based on an approximate intact mAb-1 monomer mass of 149 kDa and compositional information from complementary analyses. Because the fitted Gaussian populations are broad and partially overlapping, the indicated ±25 and ±50 kDa offsets represent approximate compositional relationships rather than unique molecular assignments. Other experimentally observed building blocks, including the ~74 kDa [LC][HC] species, may contribute within these broad populations without forming a separately resolved Gaussian component.

| Subfamily, color | Mass centroid $\bar{m}$ (kDa) | Charge centroid $\bar{z}$ | Detected-ion abundance (%) | Tentative assignment |
| --- | --- | --- | --- | --- |
| S1, dark blue | 302.49±15.75 | 31.72±2.09 | 2.6 | Dimer |
| S2, light blue | 444.61±44.57 | 38.30±2.01 | 7.0 | Trimer |
| S3, orange | 543.24±54.53 | 43.12±1.44 | 11.1 | Tetramer- ~50kDa |
| S4, Coral | 596.83±33.24 | 47.87±1.60 | 12.9 | Tetramer |
| S5, green | 695.67±37.70 | 52.50±1.31 | 9.1 | Pentamer- ~50kDa |
| S6, pale green | 729.94±39.96 | 48.80±1.73 | 7.4 | Pentamer-~25kDa |
| S7, red | 797.45±36.97 | 56.29±1.46 | 7.7 | Pentamer+ ~50kDa |
| S8, Salmon | 851.72±51.64 | 52.56±1.81 | 5.0 | Hexamer- ~50kDa |
| S9, Lavender | 913.76±43.19 | 59.63±2.06 | 8.6 | Hexamer |
| S10, Thistle | 1029.89±47.93 | 65.28±2.13 | 5.5 | Heptamer- ~25kDa |
| S11, brown | 1085.94±55.24 | 59.21±2.78 | 4.1 | Heptamer+ ~25kDa |
| S12, light brown | 1200.41±49.39 | 66.67±3.09 | 4.6 | Octamer |
| S13, magenta | 1298.42±63.30 | 74.25±2.05 | 3.2 | Nonamer- ~50kDa |
| S14, pink | 1371.52±55.02 | 68.76±2.44 | 3.0 | Nonamer+ ~25kDa |
| S15, grey | 1520.54±68.32 | 77.82±3.36 | 3.8 | Decamer+ ~25kDa |
| S16, light grey | 1731.28±79.59 | 84.63±3.18 | 2.5 | Dodecamer- ~50kDa |
| S17, yellow green | 1954.41±112.85 | 93.12±3.13 | 1.3 | Tridecamer |

Table S9. Summary of the nine-component residual-GMM model selected by BIC for the H1 fraction. Gaussian components were fitted in the $\left( \ln m , r_{z} \right)$space, where $r_{z}=\ln z-\left( a+b\ln m \right)$. The components represent statistical ion families and should not be interpreted directly as discrete molecular species. $z_{\mathrm{obs}}/z_{\mathrm{pred}}$ denotes the observed-to-predicted charge ratio, with values above or below 1 indicating relative overcharging or undercharging, respectively, after accounting for molecular mass.

| Component | Weight  percentage | Mass center | $r_{z}$center | $z_{\mathrm{obs}}/z_{\mathrm{pred}}$ |
| --- | --- | --- | --- | --- |
| C1 | 2.42% | 301.5 kDa | −0.0083 | 0.992 |
| C2 | 18.13% | 585.7 kDa | −0.0046 | 0.995 |
| C3 | 17.24% | 611.9 kDa | +0.0553 | 1.057 |
| C4 | 13.29% | 673.2 kDa | −0.0694 | 0.933 |
| C5 (excluded) | 0.74% | 815.4 kDa | +0.2632 | 1.301 |
| C6 | 12.47% | 843.3 kDa | −0.0205 | 0.980 |
| C7 | 15.48% | 884.1 kDa | +0.0430 | 1.044 |
| C8 | 10.41% | 1235.6 kDa | −0.0656 | 0.937 |
| C9 | 9.83% | 1482.8 kDa | +0.0154 | 1.016 |
